# Esophageal adenocarcinoma relapse after chemoradiation is dominated by a basal-like subtype

**DOI:** 10.1101/2025.01.28.635332

**Authors:** Sascha Hoppe, Soha Noseir, Ali Yazbeck, Nestor Zaburannyi, Jan Großbach, Su Ir Lyu, Oscar Velazquez Camacho, Stefan Müller, Florian Gebauer, Vanessa Richartz, Barbara Holz, Johannes Berg, Viktor Achter, Janine Altmüller, Kerstin Becker, Christoph Arolt, Lydia Meder, Yue Zhao, Hans Schlößer, Wolfgang W. Baus, Florian Kamp, Christian Baues, Andreas Beyer, Margarete Odenthal, Alexander Quaas, Reinhard Buettner, Axel M. Hillmer

## Abstract

Neoadjuvant chemoradiation therapy (RCT) is a frequently used treatment regimen for esophageal adenocarcinoma (EAC); however, the response varies dramatically, and resistance is a clinical challenge. We aimed to identify the molecular mechanisms underlying RCT resistance. We established a mouse xenograft RCT model with human EAC cell lines representing different response groups, and tested enhanced genomic instability as a potential evolutionary modulator by reducing *BRCA2* function. Xenografts that relapsed after RCT displayed upregulation of stress response keratins, including *KRT6* and *KRT16* connected with a basal-like transcriptomic/ proteomic phenotype. We screened our cohort of 728 patients with EAC and found significantly shorter overall survival for patients with KRT6-high tumors, driven by patients receiving neoadjuvant treatment. Overall, we identified a basal-like cell state in EAC that reflects RCT relapse. The basal-like subtype is a marker of treatment failure, providing a new avenue for translational research to overcome RCT resistance.

## Introduction

Esophageal cancer is the seventh most common cause of cancer-related deaths world wide ^1^ with esophageal adenocarcinoma (EAC) being the most prevalent subtype in Western countries. The age-adjusted incidence rate of EAC increased by more than 75% among non-Hispanic whites in the United States from 1992 to 2016 ^2^. EAC has an overall five-year survival rate of only 20% and a median overall survival of less than a year ^3^. The poor prognosis is due to the detection of the disease usually at a late stage and the lack of targeted therapies. Therefore, perioperative chemotherapy or neoadjuvant chemoradiation therapy (RCT) is the standard of care for locally advanced EAC and is used in the majority of patients with locally advanced disease ^4^. The response to treatment varies substantially, ranging from a complete pathological response to a minor or no response. Furthermore, disease recurrence with local or distal metastases ranges from 30% to >50% ^5,6^, indicating that primary and emerging resistance are major problems in the clinical care of patients with EAC. Approximately one-third of patients with EAC can be classified as minor responders to RCT and suffer from side effects of the treatment without desired clinical improvement ^7^. Therefore, biomarkers are urgently needed to spare these patients from adverse effects, underscoring the need for a better understanding of the resistance mechanisms, and to implement new treatment strategies for non-responding/early relapsing patients.

At the genomic level, EAC is a tumor type with the highest burden of small somatic mutations (single nucleotide variants [SNVs], double base substitutions, short insertions/deletions [indels], ^8^). It has the highest rate of mobile element insertions in solid tumors ^9^ and is characterized by a high level of somatic copy number alterations (SCNAs, ^10^). With these features, EAC is a prime example for cancers with extremely altered genomes. While a mutation signature associated with DNA damage response, also described as BRCAness, has been identified as the dominant signature in 20% of EAC ^10^, mutations in DNA double-strand break response genes are infrequent ^11^. However, the high level of genomic rearrangement indicates that genomic instability is an important factor in the tumor evolution of EAC. Based on the high mutation and rearrangement rates, most tumors have genomically altered cancer driver genes, including receptor tyrosine kinases such as *ERBB2, EGFR, MET*, and genes of the MAPK and PI3K pathway ^10–13^ but the response rates in clinical trials are low ^14–17^. Most likely, the low response rate is due to the complex nature of the tumor genomes, where multiple cancer driver events co-exist ^10,18^. *TP53* is the most frequently mutated gene in EAC, with a mutation frequency of 72% in the largest genome-wide analysis ^11^ underscoring genomic instability is a key feature of EAC.

In the present study, we aimed to identify the molecular characteristics of RCT resistance in EAC. We hypothesized that relapsed EAC after the initial response to RCT treatment contains features of resistance and that these features can be identified when comparing post-RCT (relapsed) tumors with those before or without RCT. Therefore, we established a xenograft mouse model using human EAC cell lines mimicking the RCT protocol for patients with EAC according to the CROSS trial ^19^. We used human EAC cell lines with *TP53* mutations to reflect genomically instable EAC. To futher increase genomic instability, the putative engine of evolution towards resistance, we genomically edited *BRCA2* to reduce the rate of homology-based DNA double strand repair in a parallel series of experiments. Comparison of sensitive vs. resistant xenograft models before and after RCT led to a transcriptomic signature of RCT resistance with remarkable similarity to induced *BRCA2* dysfunction. Systematic transcriptomic and proteomic analyses have revealed a basal-like subtype of resistant/ post RCT tumors. Using KRT6 as a marker for the basal-like phenotype in our cohort of 728 EAC patients, we found a striking association between high expression of KRT6 and short overall survival suggesting that a basal-like subtype in EAC is a predictor, and potentially a contributing factor, for RCT resistance.

## Results

### Three EAC models show response to RCT treatment protocol

To determine the impact of RCT treatment on the genomic evolution of EAC tumors, we chose three representative EAC cell lines, Eso26, OE19, and OE33x (OE33 recultured after subcutaneous xenograft), which have been reported to form tumors in mice ^20^. We established an RCT protocol to mimic the treatment regime applied to patients with EAC according to the CROSS protocol ^21^. First, we tested the sensitivity of these cell lines to RCT treatment *in vitro*. The RCT treatment involved varying concentrations of two chemotherapeutic agents, Carboplatin and Paclitaxel, combined with a 2 Gy radiation dose. The three cell lines showed different responses to the RCT treatment. OE19 exhibited the least sensitivity, necessitating a 100 nM of Carboplatin and 10 nM of Paclitaxel IC50 dose, while OE33x displayed a higher sensitivity with 1 nM of Carboplatin and 0.1 nM of Paclitaxel IC50 dose. Notably, Eso26 emerged as the most responsive cell line, showing remarkable sensitivity at low dose of 0.1 nM Carboplatin and 0.01 nM Paclitaxel (**Fig. 1A**). Second, the tumor growth potential of the three models was observed *in vivo* as the cell lines were injected into immunocompromised nude mice, and the growth rates of xenograft tumors were monitored (**Fig. 1B-D** and **Supplementary Fig. 1A-C**). Remarkably, Eso26 exhibited the fastest pretreatment growth rate, reaching tumor sizes greater than 500 mm³ in 20 days, followed by OE19, which reached approximately 200 mm³ in the same period. In contrast, OE33x displayed slower growth, requiring twice the time (up to 40 days) to reach approximately 100 mm³ (**Fig. 1D**, and **Supplementary Fig. 1A-C**). This divergence in sensitivity and growth behavior across the three EAC cell lines allowed us to explore the nuanced evolution of genomic dynamics under the effect of RCT treatment. In summary, three EAC cell lines with different RCT sensitivity were established as xenograft models.

**Figure 1:**
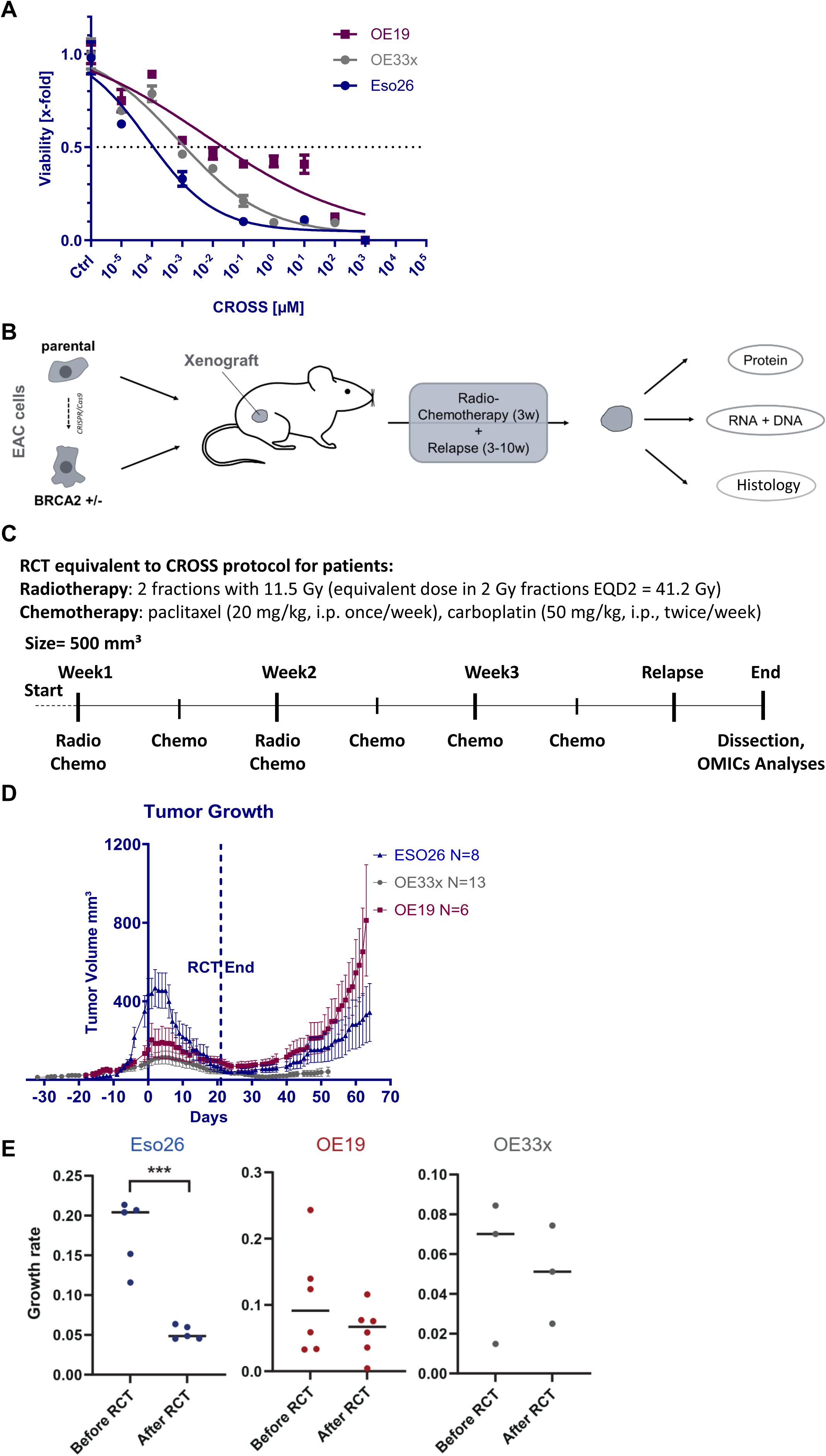
RCT treatment response tested in three EAC models. **(A)** Cell Titer Glo (CTG) viability assay illustrates the distinct responses of Eso26, OE19, and OEx33 esophageal adenocarcinoma (EAC) cell lines to RCT treatment. CROSS chemotherapeutic concentrations displayed on the x-axis are calculated as µM Carboplatin and µM*10 Paclitaxel concentrations combined with 2 Gy of radiation. **(B)** Experimental design and workflow for *in vivo*-based analyses. **(C)** Overview of the combined RCT treatment protocol executed *in vivo*. **(D)** Graphical representation of growth curves for the three EAC tumor models, showcasing the initial response to RCT treatment and subsequent post-treatment progression in 4 out of 13 tumors in the OE33x model, 4 out of 6 tumors in the OE19 model, and in 6 of 8 tumors in the Eso26 model following RCT treatment. FFPE refers to formalin-fixed, paraffin-embedded tissue; N, numbers of tumors included in the analysis. **(E)** Growth rates of three EAC xenograft models before and at day 40 after RCT start. ***, P < 0.001 T-test.

### RCT treatment induces differential patterns of tumor progression *in vivo*

The *in vivo* experimental workflow followed the scheme outlined in (**Fig. 1B, C**). We examined the treatment responses of the selected models to the CROSS protocol. RCT treatment was ideally initiated at 500 mm³ (day 0), but the differences in the tumor sizes at treatment initiation (500 mm³ for Eso26, 200 mm³ for OE19, and 100 mm³ for OE33) were due to distinct initial growth rates between the models. The three tumor models underwent RCT treatment for three consecutive weeks, concluding on day 21. Radiation was applied locally to the tumor regions in two fractions of 11.5 Gy each with 10 MeV electrons (week one and week two) resulting in an equivalent dose in 2 Gy fractions of 41.2 Gy (assuming αβ = 10). Subsequently, the progression patterns of the treated tumors were monitored for an additional 3-6 weeks (in total 42-63 days from start of the treatment). Our data revealed that tumors progressed after the termination of RCT treatment, and most of them reached a sufficient size, enabling subsequent omics analysis. Interestingly, OE19 tumors showed a higher progression rate post-RCT than Eso26, reaching ∼1000 mm³ on day 63, after treatment termination, in agreement with the poor response *in vitro*. Eso26 tumors exhibited a growth rate half as fast as OE19, reaching ∼500 mm³ after 63 days, indicating that RCT enhanced potential resistance in fast-growing OE19 tumors (**Fig. 1D**). The growth rate of Eso26 xenografts slowed down significantly after RCT (P = 0.0002, T-test, **Fig. 1E**), suggesting a substantial adaptation to therapy to reach resistance. As anticipated, the slow-growing OE33x tumors showed the slowest recovery, as the growth rate further decelerated after RCT treatment. It is important to highlight that the individual tumor growth curves demonstrated a prevalent pattern of progression (i.e., tumor regrowth after regression), as observed in 6 out of 8 tumors in the Eso26 model and similar patterns in OE19 and OE33x (**Supplementary Fig. 1A-C**). Overall, the *in vivo* RCT responses were in agreement with the *in vitro* behavior, reflecting different EAC response types.

### *BRCA2* is essential for EAC models and reduced levels lead to slow tumor growth

Since EAC is characterized by genomic instability measured by a large number of somatic copy number alterations (SCNAs) ^13^, and extreme genomic instability in EAC can be driven by somatic *BRCA2* mutations ^10^, we investigated whether the reduced function of *BRCA2* could accelerate genomic changes leading to RCT resistance. We used CRISPR/Cas9 to knockout *BRCA2* and analyzed 120 clones across the three EAC cell models. While almost all clones showed genomic editing of *BRCA2*, none of the clones showed frameshift mutations of all alleles (the Eso26 cell line has three copies, OE33 >5 copies ^22^, OE19 three copies [own data] of the *BRCA2* locus, **Fig. 2A** and **Methods**). We concluded that *BRCA2* knockout is lethal in the three EAC cell models, underscoring the pivotal molecular role of *BRCA2* in EAC cells. Instead, we opted for clones with a knockout of all but one allele (e.g., knockout of two out of three alleles), referred to hereafter as *BRCA2* knockdown (*BRCA2*kd). As a result of the reduction in *BRCA2* expression, the protein level of BRCA2 was significantly downregulated in *BRCA2*kd cells compared with their parental counterparts (**Fig. 2B**).

**Figure 2:**
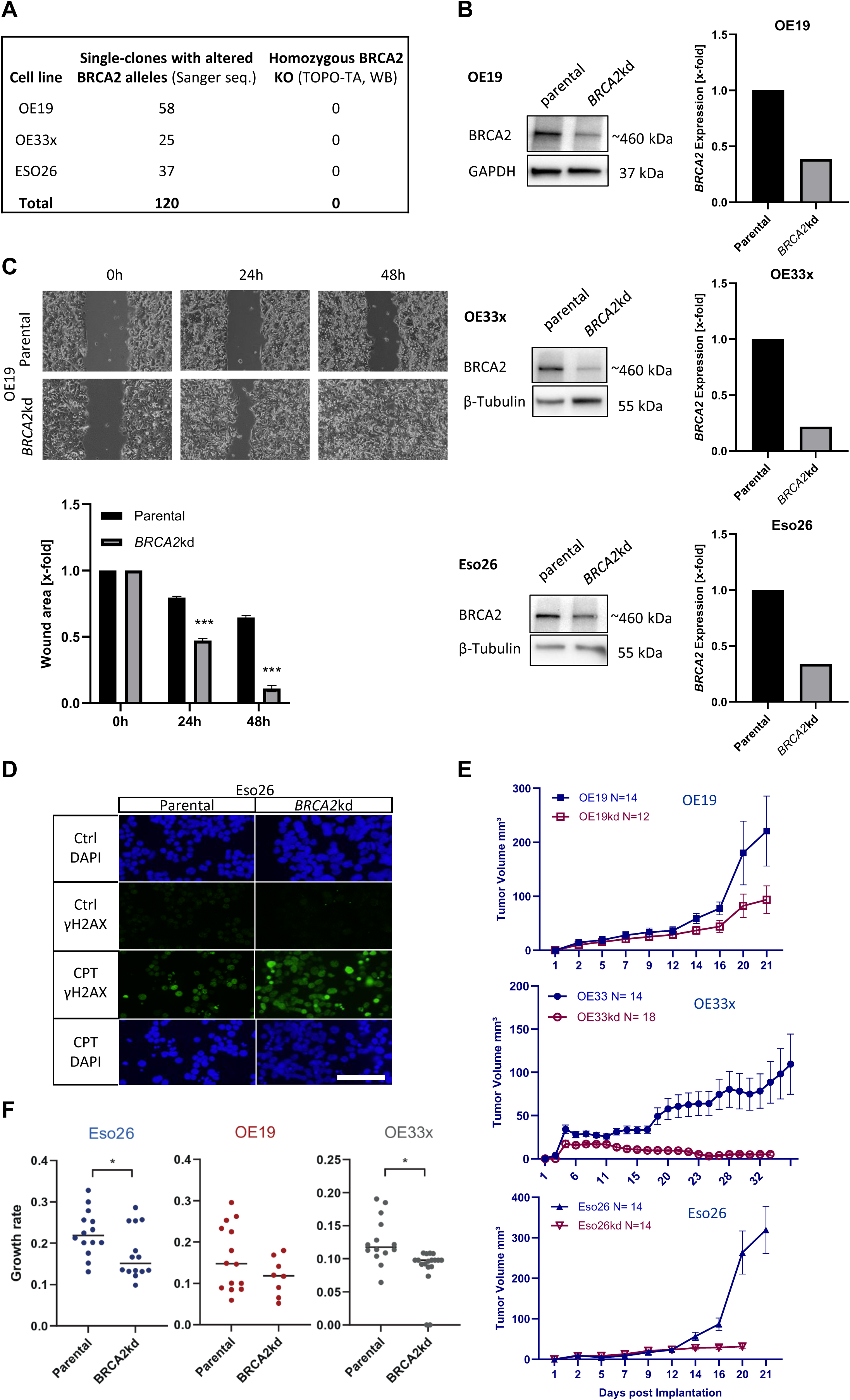
Reduced function of *BRCA2* leads to changes in migration potential and DNA damage response in EAC tumor cell models. **(A)** CRISPR/Cas9 mediated *BRCA2* allele alteration was induced in 120 clones across three EAC cell lines. The resulting clones were verified through Sanger sequencing. Notably, none of the viable *BRCA2* edited clones had a homozygous *BRCA2* knockout (KO). The *BRCA2* edited clones have one wild type/in-frame *BRCA2* allele left, referred to as *BRCA2*kd. **(B)** Verification of *BRCA2*kd across EAC models through Western blot analysis. **(C)** Top: OE19 *BRCA2*kd cells exhibit enhanced migration rates, as assessed through a wound-healing assay, in comparison to parental cells at 0, 24, and 48 hours (* P<0.05, T-test). Bottom: ImageJ analysis of inverted microscopic measurements were employed to measure the distance traveled by cells. **(D)** Immunofluorescence staining of γH2AX in Eso26 parental and *BRCA2*kd cells. The expression of the DNA damage marker γH2AX increased in 50 µM Carboplatin-treated Eso26 *BRCA2*kd tumors compared to parental tumors 40x magnification. CPT, Carboplatin. **(E)** Xenograft growth dynamics in parental and *BRCA2*kd tumors. *BRCA2*kd significantly decreased the tumor growth potential *in vivo* when compared to parental tumors. **(F)** Growth rates of three EAC parental and *BRCA2*kd xenograft models. *, P<0.05 T-test.

Next, we investigated the effect of the reduction in *BRCA2* function on EAC cells. *BRCA2*kd cells exhibited significant gains in migration capabilities compared to the parental cells across the three cell lines (**Fig. 2C** and **Supplementary Fig. 1D and E).** Additionally, we observed a notable increase in the expression of the DNA damage marker γH2AX in Eso26 *BRCA2*kd cells treated with Carboplatin (**Fig. 2D**), underscoring the impact of the *BRCA2*kd phenotype on DNA damage response in EAC cells. Furthermore, we explored the influence of *BRCA2*kd on tumor growth and development *in vivo*. The results revealed that *BRCA2*kd led to a lower tumor growth rate than the parental cell line xenografts in all three tumor models (**Fig. 2E**), reaching significance for Eso26 and OE33x (P = 0.033 and P = 0.002, respectively; T-test, **Fig. 2F**). Notably, in the OE33x model, we observed a substantial decrease in *BRCA2*kd tumor size over 35 days post-implantation, and no tumors were observed afterwards. Moreover, we investigated the response of *BRCA2*kd tumors to RCT treatment. Eso26 *BRCA2*kd tumors exhibited distinctly smaller tumor volumes than OE19 *BRCA2*kd tumors, which displayed greater tolerance to such alterations at the start of RCT treatment. Notably, OE19 *BRCA2*kd tumors regrew at a fast rate comparable to their *BRCA2* proficient counterparts, reflecting rapid recovery (**Supplementary Fig. 1F**). In contrast, Eso26 *BRCA2*kd tumors showed consistently low tumor growth rates, indicating more pronounced suppression of progression (**Supplementary Fig. 1G**).These data highlight the critical role of *BRCA2* in DNA repair, cell proliferation, and EAC tumor growth.

### *BRCA2*kd and RCT treatment induce marked genomic changes in EAC tumors

Next, we investigated whether genomic changes induced by *BRCA2*kd alone could lead to genomic instability, how instability compared with changes induced by RCT treatment, and whether *BRCA2*kd accelerates RCT-induced changes. Accordingly, we performed whole exome sequencing (WES) and examined the difference in the genomic instability index (GII) between the Eso26 and OE19 parental/*BRCA2*kd tumor models that underwent RCT (no tumor growth for OE33 *BRCA2k*d). GII was induced at a higher level in *BRCA2*kd tumors compared to RCT treatment in the parental, *BRCA2* proficient counterparts **(Fig. 3A; Supplementary Fig. 2A; Supplementary Table 1)**. Notably, RCT treatment of *BRCA2*kd tumors did not result in additional increases, but rather equal GII compared to non-treated *BRCA2*kd tumors in both Eso26 and OE19 models, suggesting that a maximum tolerable genomic instability was reached. In addition, we investigated the characteristics of acquired SNVs and indels. When comparing post-RCT and *BRCA2*kd xenografts with their parental non-treated, non-edited counterparts, we observed no statistically significant differences in mutation numbers when comparing the effect. Furthermore, we conducted mutational signature analysis in response to RCT and *BRCA2*kd. We identified mutation signatures SBS1 (aging), SBS3 (BRCA deficiency), and SBS5 (unknown etiology, clock-like, increased in bladder cancer with *ERCC2* mutations, and in cancers due to tobacco smoking; **Fig. 3B**). RCT, *BRCA2*kd, and *BRCA2*kd+RCT did not show consistent differences across xenograft models with regard to the contribution of individual signatures (**Supplementary Fig. 2B**). Overall, the genomic profiles may indicate that reduced *BRCA2* function results in a maximum of tolerable genomic instability.

**Figure 3:**
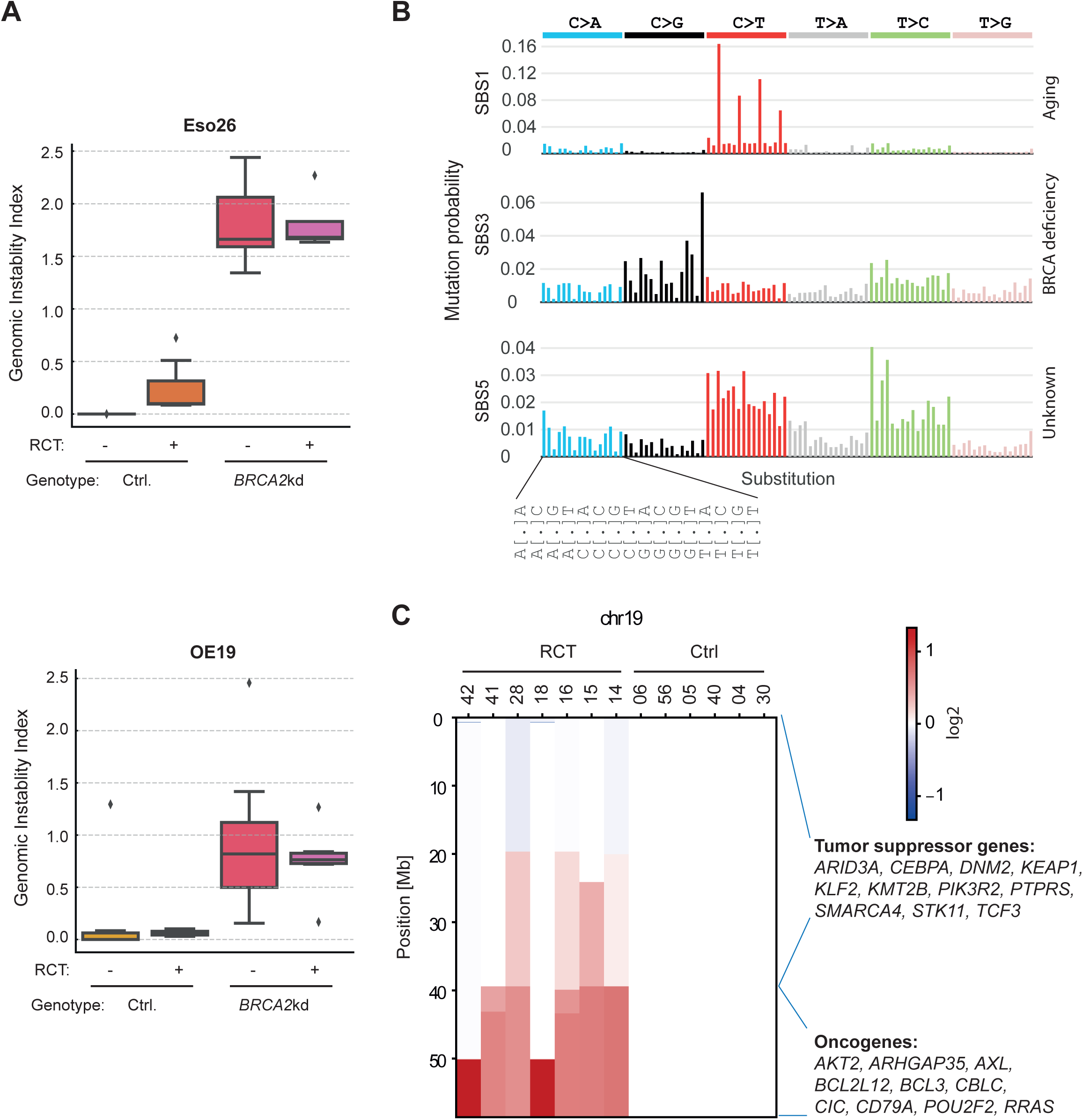
Induced genomic response in EAC parental and *BRCA2*kd tumors post RCT treatment. **(A)** Genomic copy number analysis based on WES defines the genomic instability index (GII, y-axis) as a measure for the fraction of the genome that is altered. RCT treatment induces GII to a moderate extent while *BRCA2*kd leads to high GII in Eso26 and OE19 tumor models. **(B)** Single nucleotide variant (SNV) analysis of mutations acquired during the course of the experiment compared to parental cell lines without treatment reveals three mutation signatures. **(C)** Copy number profile of chromosome 19 (y-axis) of post-RCT relapsed Eso26 tumors and parental Eso26 as reference is shown with red indicating gained copy number. Note consistent copy number gain of chromosome 19q in Eso26 RCT-treated tumors compared to treatment-naive tumors. Known cancer-related genes are shown on the right.

Next, we aimed to identify recurrent genomic events that could explain the development of resistance after RCT. At the level of acquired SNVs and indels during the experiments, we found that non-silent mutations were enriched in genes listed in the Cancer Gene Census ^23^ or OncoKB ^24^ (p < 0.0001, χ^2^ Test, both gene categories), including non-silent mutations in *ATM, BRCA1, KDM5C, MUC4, NOTCH1, TERT,* and *MAP3K13* (**Supplementary Table 2**).

However, many of these mutations showed low allelic fractions, and none of them were classical cancer driver mutations that could explain relapse after RCT. This implies that the accumulation of small effects, rather than a single strong mechanism, might contribute to relapse. Interestingly, somatic copy number alteration (SCNA) data analysis identified a copy number gain on chromosome 19q in all six Eso26 RCT-treated compared to treatment-naive tumors, highlighting the presence of potential candidate genes for a post-RCT phenotype or the development of resistance **(Fig. 3C)**. The genomic region contains oncogenes *AKT2, AXL, BCL2L12, BCL3, CBLC, CIC, CD79A, POU2F2, RRAS,* and *ARHGAP35* as potential supporters of RCT resistance. Of note, *BCL2L12* was the only oncogene within the copy gain region of xenografts Eso26-RCT-42 and -18, with the smallest common copy gain regions (**Fig. 3C**). Overall, *BRCA2*kd results in expected genomic instability and RCT does not lead to the emergence of small mutations in recurrent driver genes, and 19q copy gain in the sensitive Eso26 model correlates with relapse.

### Transcriptomic and proteomic profile changes in *BRCA2*kd tumor models exceed changes in tumors after RCT treatment

To understand the nature of biological changes induced by *BRCA2*kd and RCT, we performed transcriptomic and proteomic analyses. First, we examined sample similarity across the three chosen tumor models by employing the Euclidean distance matrix based on both omics approaches. The results indicated that the molecular signatures were dominated by EAC cell line types, reflecting the importance of inter-patient differences (**Fig. 4, Supplementary Tables 3, 4**). The second strongest factor influencing overall transcriptomic and proteomic profiles was *BRCA2*kd. Unexpectedly, the *BRCA2*kd signature was more dominant than the RCT signature, defined by shorter Euclidean distances. However, RCT also resulted, although to a lesser extent, in transcriptomic and proteomic signatures that led to the separation of RCT and non-RCT tumors in most samples.

**Figure 4:**
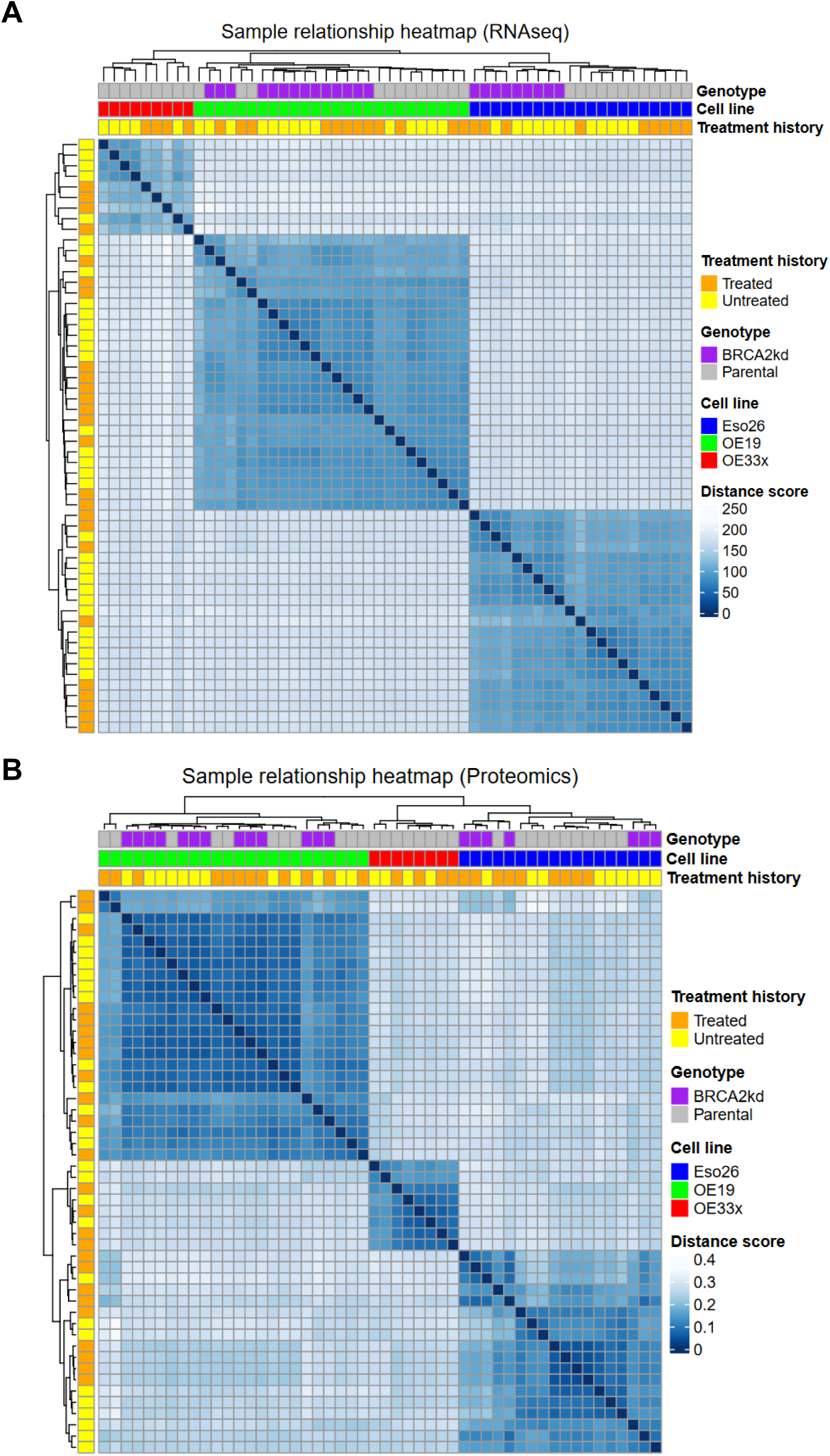
Transcriptomic and proteomic profile changes are most prominent in *BRCA2*kd tumor models. **(A)** Hierarchical clustering of the Euclidean distance matrix based on 3’ mRNASeq normalized gene expression of the three EAC tumor models. This heatmap showing sample similarity and the dominant signature of EAC cell lines. **(B)** Hierarchical clustering of the Euclidean distance matrix based on proteomic data of the three EAC tumor models. Heatmap illustrates proteins that are differentially expressed across all groups in the EAC tumor models.

### *BRCA2*kd and RCT treatment induce overlapping transcriptomic changes in EAC tumor models

We conducted an extensive analysis of the transcriptomic profile changes after *BRCA2*kd and RCT treatment. Our data revealed that the highest number of differentially expressed genes (DEGs) was induced in *BRCA2*kd of the Eso26 tumor model, with 2,986 DEGs compared to parental tumors (**Fig. 5A, Supplementary Fig. 3**). We also observed a substantial change in 586 DEGs in Eso26 RCT-treated tumors compared to their non-treated parental counterparts. In contrast, the transcriptomic analysis of OE19 RCT-treated tumors displayed the least changes, with only 10 DEGs compared to their parental counterparts, whereas OE33 showed an intermediate number of DEG in the RCT/parental comparison (**Fig. 5A**). To investigate whether the RCT-induced expression changes in parental Eso26 (model with the strongest RCT expression differences) showed similarities with the expression differences between parental Eso26 and OE19 (least RCT expression differences and most primary resistant model), we systematically compared the expression differences between the two comparisons. We observed a significant correlation between the fold-expression changes across the two comparisons (r = 0.3799, p < 0.0001, Pearson correlation test, **Fig. 5B**), supporting the assumption that the gene set is resistance-associated. Since the Eso26 *BRCA2*kd model showed fewer transcriptomic changes upon RCT than the parental Eso26 (**Fig. 5A**), it seems possible that *BRCA2*kd shifted EAC to a more resistant phenotype. Therefore, we tested for correlation between RCT- and *BRCA2*kd-induced changes in expression. Indeed, we observed a strong correlation between the expression changes in the parental Eso26 / RCT and parental Eso26 / *BRCA2*kd comparisons (r = 0.6407, p < 0.0001, Pearson correlation test, **Fig. 5C**), suggesting that *BRCA2*kd induces similar changes as RCT resistance.

**Figure 5:**
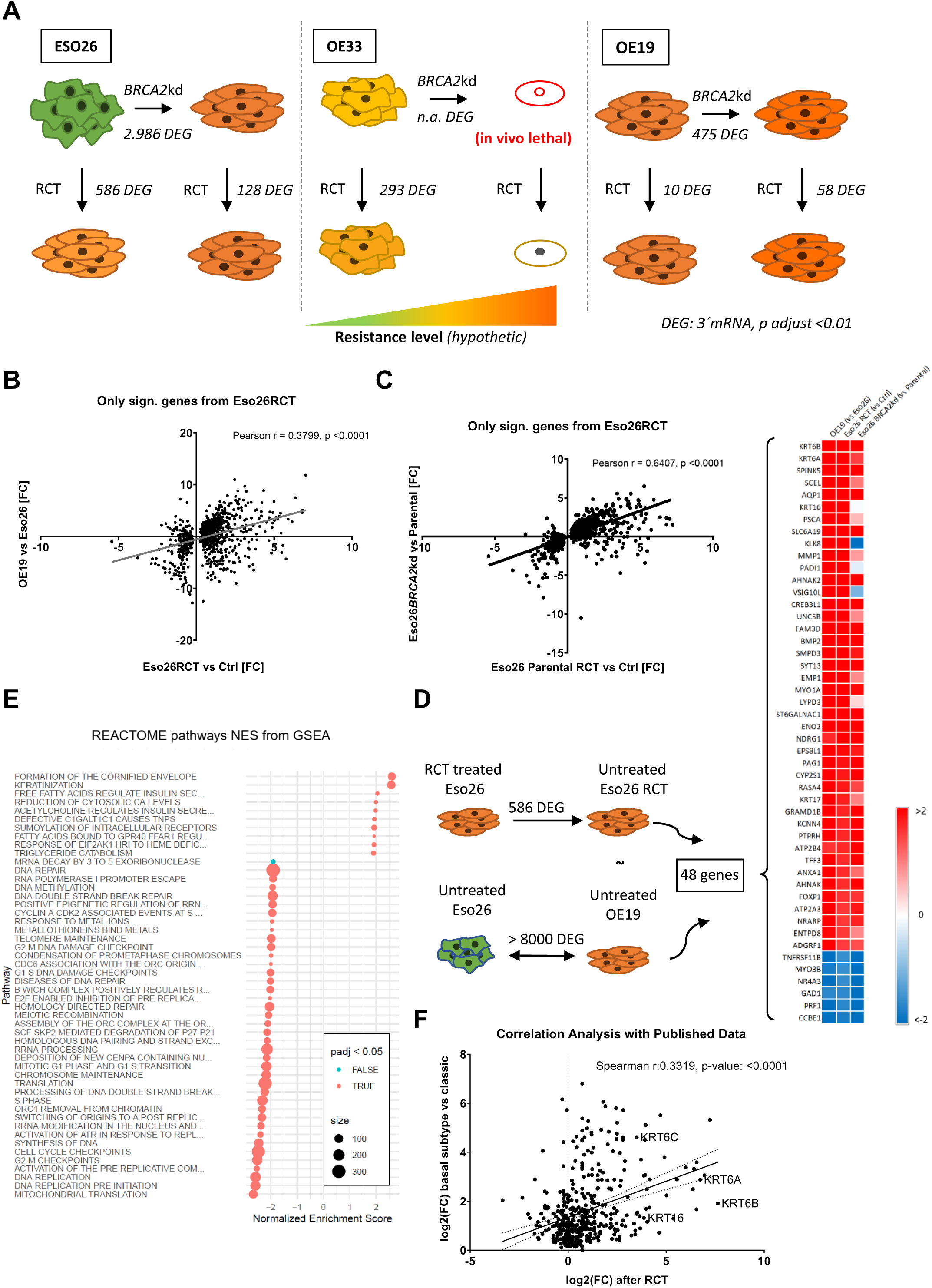
*BRCA2*kd and RCT treatment induced overlapping transcriptomic changes in EAC tumor models. **(A)** Illustration depicting the numbers of DEGs across the three EAC tumor models. **(B)** Correlation analysis of gene expression differences between RCT-treated vs. untreated Eso26 tumors (x-axis) and untreated OE19 vs. Eso26 tumors (y-axis). **(C)** Correlation analysis of gene expression differences between RCT-treated vs. untreated Eso26 tumors and untreated Eso26 BRCA2kd vs. untreated Eso26 parental tumors. **(D)** Definition of resistance expression signature based on common expression differences between Eso26 treated vs. untreated and OE19 vs. Eso26 (left schematic and left and middle column in heatmap). Resistance gene set of 48 genes correlates with transcriptomic changes induced by *BRCA2*kd in Eso26 (right column in heatmap). **(E)** GSEA for REACTOME pathways for differentially expressed genes between Eso26 RCT vs. Eso26 parental. **(F)** Correlation analysis between gene expression differences of RCT treated Eso26 vs. untreated tumors and the basal-like keratinization vs. classic subtype reported by Guo et al. 2018 ^25^.

To derive a robust EAC expression gene set for RCT resistance, we extracted overlapping DEGs from the two comparisons with resistant xenografts (Eso26 RCT vs. Eso26 Ctrl. and OE19 Ctrl. vs. Eso26 Ctrl.), excluded weakly expressed genes, and focused on genes with RefSeq IDs (**Supplementary Table 5**). The filter resulted in 48 genes (DEGs) as RCT resistance signature including upregulation of several keratins (*KRT6, KRT16*, and *KRT17*)*, MMP2, BMP2, NDRG1, TFF3,* and *FOXP1* (**Fig. 5D**). Interestingly, we observed a high density of signature genes on 19q (**Supplementary Fig. 2C**), suggesting that 19q gain contributed to the RCT signature in Eso26. Next, we aimed to better understand of the pathway changes behind resistance-associated expression changes and performed gene set enrichment analysis (GSEA) for Eso26 tumors after RCT treatment compared with their untreated counterparts. The data showed that common downregulated pathways were related to proliferation, including DNA replication and G2/M/cell cycle checkpoints, whereas common upregulated pathways were related to cornification/keratinization (**Fig. 5E**). The latter involves several keratins as markers of basal cells and epithelial differentiation.

A privious study reported that there are two distinct EAC transcriptome subtypes: basal and classic. The basal-like subtype is reportedly more resistant to chemotherapy than the classic subtype ^25^. By integrating the published transcriptomic data of the basal-like subtype with our data from Eso26 RCT-treated tumors, we observed a positive correlation between the two groups (r = 0.3319, p < 0.0001, Spearman’s correlation test, **Fig. 5F**). This correlation suggests a connection between the basal-like subtype, keratin upregulation, and the development of resistance to RCT treatment. We searched for transcription factors that might be responsible for the switch towards the basal-like phenotype and identified *ATF3* as a potential candidate. *ATF3* was significantly upregulated in Eso26-RCT, Eso26-*BRCA2*kd, OE19-RCT, and OE19-*BRCA2*kd-RCT, and was more highly expressed in the resistant OE19 cell line than in the sensitive Eso26 cell line (**Supplementary Table 5**). *ATF3* is a stress-induced transcription factor that characterizes quiescent colorectal cancer cells ^26^.

In summary, we have defined a transcriptomic signature for a post-RCT/ RCT-resistance phenotype. This resistance phenotype is similar to the changes induced by *BRCA2*kd, and has basal-like characteristics.

### Elevated expression of the basal-like cell marker keratin 6 is associated with short survival in neoadjuvant-treated EAC patients

To investigate the potential role of keratinization in the development of RCT resistance, we examined the overexpression of keratins in our dataset of RCT-treated tumors. Our transcriptomic data revealed that several keratins, including *KRT6, KRT16,* and *KRT17*, were upregulated in RCT-treated Eso26 tumors (**Fig. 6A**). Cytokeratin 5/6 (CK5 and CK6 encoded by *KRT5* and *KRT6*) are markers for squamous cell carcinomas of basal cell origin. Both keratins occur in the squamous epithelium, KRT5 dominates in basal and KRT6 in suprabasal layers ^27,28^. We therefore used KRT6 as a marker for the transdifferentiation to a basal-like transcriptomic phenotype. We validated the upregulation of KRT6 upon RCT through immunohistochemical (IHC) staining of the xenografts, demonstrating KRT6 overexpression in RCT-treated tumors (**Fig. 6B**). To evaluate the clinical significance of these findings, we analyzed 1,161 EACs collected before and after neoadjuvant treatment, respectively, from patients who underwent surgical resection. The histological examination of EAC tumors from treated and untreated patients revealed varying levels of KRT6 expression in the 728 tumors that could be evaluated, classified as 0 (no expression, n = 541, 74.3%), low (expression in <25% tumor cells, n = 117, 16.1%), and high (expression in >25% tumor cells, n = 70, 9.6%; **Fig. 6C, D**). The overall survival analysis of the 728 patient cohort showed significantly shorter survival for patients with KRT6 high-expressing tumors (score 2, p = 0.0054, log-rank test, hazard ratio [HR] = 1.2, **Fig. 6E**). Strikingly, when we separated the cohort into patients with and without neoadjuvant treatment (n = 497 and n = 231, respectively), we found a statistically significant shorter survival only for patients with KRT6 high-expressing tumors in the neoadjuvant-treated group (p = 0.00071, log-rank test, HR = 1.2). This data suggests that the basal subtype marker KRT6 might be used as a biomarker for poor response to neoadjuvant treatment in EAC patients.

**Figure 6:**
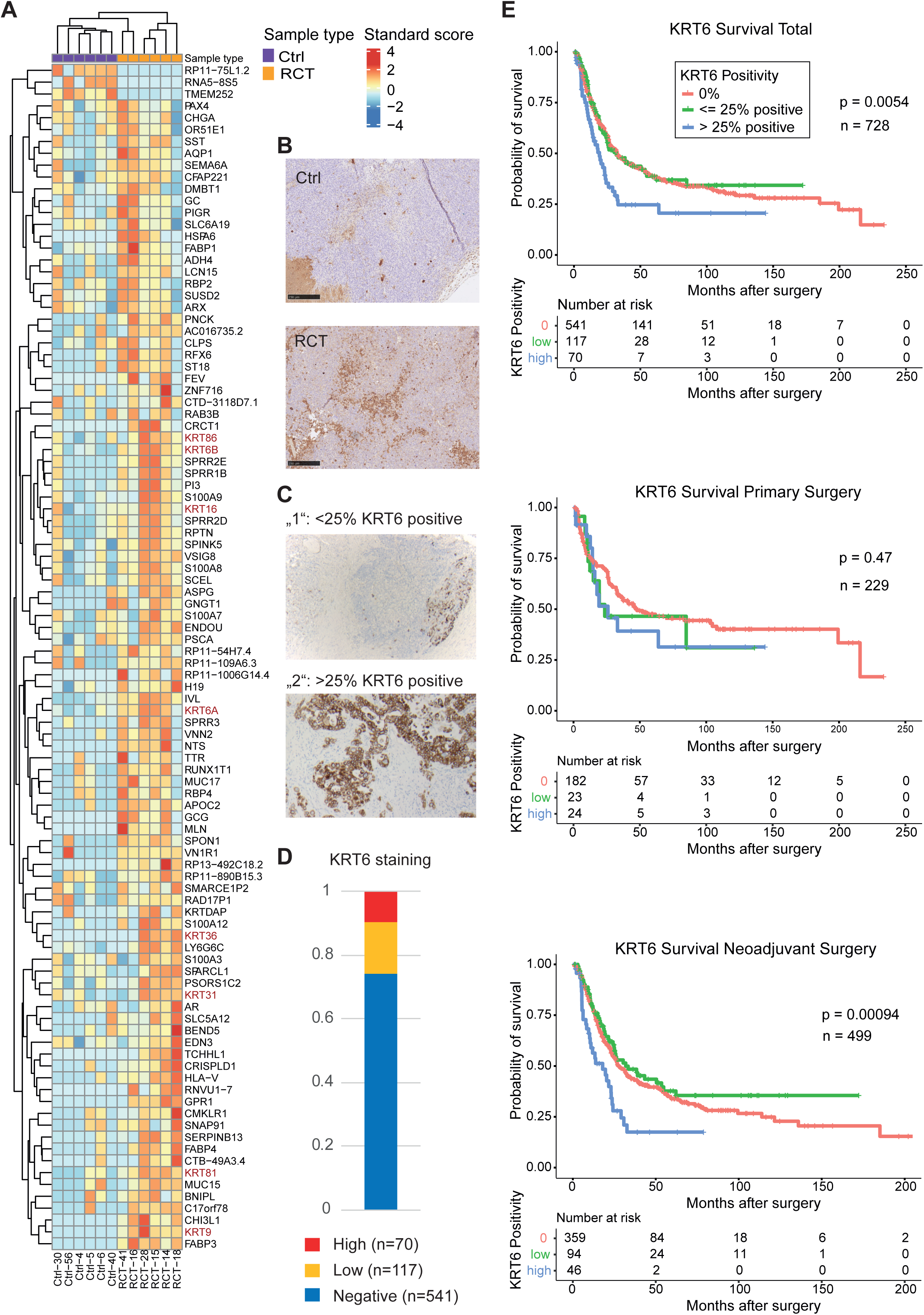
Elevated expression of basal cell marker keratin 6 (KRT6) is associated with poor survival in EAC patients who underwent neoadjuvant treatment. **(A)** Gene expression heatmap shows the overexpression of several keratins in RCT-treated Eso26 tumors. **(B)** Immunohistochemical (IHC) staining of KRT6 in a tumor with (RCT) and without treatment (Ctrl) shows overexpression of KRT6 in treated Eso26 tumors. **(C)** Examples for KRT6 expression levels low and high for the analysis of tissue microarray (TMA) by IHC showing the differential expression levels of KRT6 in EAC patient tumors. . **(D)** Quantitative analysis of KRT6 IHC of 1161 EACs from Cologne arranged as TMA. Zero represents KRT6 negative, low indicates <25% KRT6 positive, and high signifies >25% KRT6 positive tumor cells. **(E)** Kaplan-Meier survival analysis of 728 EAC patients with evaluable IHC and survival data with different levels of KRT6 expression. Analysis of all patients is shown on top (HR = 1.201; 95%CI = 1.032 - 1.397), treatment naive patients (middle; HR = 1.189; 95%CI = 0.8977 - 1.574), and EAC of patients post-treatment at the bottom (HR = 1.202; 95%CI = 1.002 - 1.442).

To further characterize the cell state changes captured by KRT6, we used the transcriptomic data of all xenografts across all cell lines and experimental conditions and correlated gene expressions with KRT6A and KRT6B, respectively, both recognized by our KRT6 antibody. We observed a significant positive correlation between *KRT6A* and *KRT6B* (Spearman r = 0.44, p = 0.00077, **Supplementary Fig. 4**). *KRT6A*-correlated genes showed enrichment for gene ontology terms of hair follicle development, hemidesmosome assembly, and epidermis morphogenesis, while KRT6B-correlated genes were enriched for blood vessel endothelia and response to oxygen radicals (**Supplementary Fig. 5, Supplementary Table 6**). Negatively correlated with *KRT6A* were Wnt signaling and DNA replication, whereas *KRT6B* was negatively correlated with interferon-gamma-mediated signaling, type I interferon signaling, and DNA replication. The correlations suggest that KRT6-positive EAC xenografts are less proliferative, immune silent, and have a changed epithelial phenotype dealing with reactive oxygen species.

Taken together, KRT6, a marker of a resistance-associated basal-like subtype, is distinctly expressed in 25% of the tested EAC cohort and serves as a strong predictor of poor survival, particularly among patients treated with RCT.

## Discussion

An advantage of *in vitro* and mouse models is the possibility to repeatedly challenge genetically homogeneous cancer cells or tumors with the same therapy to test for recurrent molecular features that occur in response to developing resistance or relapse. Using patient-derived cancer cell lines, the genomes/transcriptomes/proteomes of experimental biological replicates can be considered almost equal at the start of our experiments. Deep omics analyses of such xenografts after RCT therefore have a higher chance identifying recurrent resistance mechanisms among a small number of tumors compared to analyses of tumors across a cohort of patients who typically have a high inter-patient genomic variability. We have applied this approach using three EAC cell lines with different sensitivities to RCT and their derivatives with *BRCA2*kd. We observed remarkable transcriptomic changes after knocking down *BRCA2* which were stronger than the long-term influence of RCT (**Fig. 4A**). *BRCA2*kd resulted in some expected phenotypes, i.e., γH2Ax as a marker for DNA double-strand breaks was increased after Carboplatin treatment (**Fig. 2D**). *BRCA2*kd tumors exhibited higher genomic instability compared to RCT-treated parental tumors in both the Eso26 and OE19 models (**Fig. 3A**). We observed an increased cell migration after knockdown of *BRCA2* for all three cell lines (**Fig. 2C, Supplementary Fig. 1D, E**). This effect has been described for prostate cancer cells previously, where a connection with high levels of reactive oxygen species has been found to be a consequence of *BRCA2* reduction and a prerequisite for the migration phenotype ^29^. Interestingly, *BRCA2* knock out seems to be lethal for the three EAC cell lines Eso26, OE19, and OE33. This might be based at least in part on the fact that all three cell lines have *TP53* mutations since mutual exclusivity of *TP53* and *BRCA1/2* mutations has been reported for gastric cancer ^30^.

This is in contrast to reported *BRCA2* knockout cancer cell lines, including the colorectal cancer cell line HCT116 ^31^ and cell lines derived from a breast cancer mouse model ^32^. Although we attempted to knock out *BRAC2* only in a limited number of EAC cell lines, it suggests that *BRCA2* is essential for many EACs. It is in agreement with the fact that germline *BRCA2* mutations are known to increase the risk to develop breast, ovarian, and a number of other cancers but not to a relevant extent EAC ^33,34^. Further, *BRCA2*kd resulted in a significantly reduced tumor growth rate (**Fig. 2F**), suggesting a high cost or increased stress that reduces the proliferation rate *in vivo*.

Strikingly, a region on chromosome 19q showed copy gains in all six post-RCT Eso26 xenografts (**Fig. 3D**). Eso26 is hypodiploid, having only one copy of chromosome 19 ^22^. The consistent copy gains suggest the presence of a gene in this region contributing to RCT resistance. The oncogenes *AKT2, AXL, BCL2L12, BCL3, CBLC, CIC, CD79A, POU2F2, RRAS,* and *ARHGAP35* within the 19q region are candidates for this phenomenon, of which *BCL2L12* is the only oncogene within the smallest common copy gain region of Eso26-RCT-18 and -42. While the deletion of 19q13.31–33 and within this region *LIG1* have been described as a mechanism for Carboplatin resistance in triple-negative breast cancer ^35^, and there is evidence for *BCL2L12* knockdown in breast cancer cells to inhibit Cisplatin-induced apoptosis ^36^ the apoptotic influence of *BCL2L12* is controversial, showing cell type dependency ^37^. *BCL2L12* inhibits post-mitochondrial apoptosis in glioblastoma ^38^ and has an anti-apoptotic effect in ovarian cancer ^39^. Therefore, *BCL2L12* might be a candidate for explaining the RCT-associated copy gain in the Eso26 xenografts and may contribute to RCT resistance.

On the transcriptomic and proteomic level, the molecular distance between xenografts was dominated by the parental (original) EAC cell lines (**Fig. 4**), indicating that inter-patient differences are larger than *BRCA2*kd and RCT-induced changes. *BRCA2*kd led, in particular on the transciptomic level, to the second strongest changes, followed by RCT, emphasizing the relevance of *BRCA2* function in the context of EAC. By differential expression analysis of post-treatment relapsed vs. non-treated and resistant cell line vs. sensitive, we defined a 48-gene resistance signature (**Fig. 5B, D**). Unexpectedly, the post-RCT resistance signature of Eso26 is highly correlated with transcriptomic changes induced by *BRCA2*kd (**Fig. 5C**), suggesting again that the stress responses have commonalities. The pathways that showed the strongest enrichment of overexpressed genes are ‘formation of cornified envelope’ and ‘keratinization’ (**Fig. 5E**). This was in part based on the induced expression of several keratins (**Figs. 5D** and **6A**) with *KRT6A* and *KRT16* known as a stress response in skin upon damage and UV light ^40,41^. Both keratins might be seen as markers for RCT-stressed EAC. *KRT16* has been found to be upregulated upon induced DNA damage in human lung epithelial cells along with a squamous transdifferentiation ^42^ and squamous differentiation seems to protect cells from apoptosis ^43^. The RCT-induced biological process we observe in the EAC model reflects a transdifferentiation as stress reponse.

The mechanistic chain of action of how RCT induces the transdifferentiation remains to be clarified. The stress-induced transcription factor *ATF3* is a potential candidate in this process. Its expression has been observed in quiescent colorectal cancer cells ^26^ resembling the less proliferative state of the *KRT6* high post-RCT state in our EAC model. While the effect of *ATF3* can be pro or contra cancer ^44,45^, the reported requirement of *Atf3* for a higher metastatic burden after chemotherapy in a mouse model ^46^ supports the involvement of *ATF3* in the observed RCT response in our EAC model. Of note, our analyses excluded mouse sequence-specific reads/peptides so that *ATF3* can be assigned to the cancer cells rather than the tumor microenvironment as in the study of Chang and colleagues ^46^.

EAC has been divided into ‘basal’ and ‘classical’ based on their transcriptomic profiles, where *KRT6A* and *KRT16* are among the core enrichment genes for the basal subtype ^25^. Indeed, we observed an overall transcriptomic shift after RCT of our Eso26 model towards the basal signature (**Fig. 5F**). We therefore use the term ‘therapy-induced basal-like’ to describe the EAC phenotype related to post-RCT and/or RCT resistance. The terminology ‘basal-like’ in cancer has been established for breast cancer, where it is related to but not identical with triple-negative breast cancer, a difficult-to-treat subtype without expression of estrogen and progesterone receptors and *HER2* ^47,48^. Basal-like breast cancers are characterized by high histological grade and proliferative rate ^49,50^, a high expression of genes of the basal cell layer of the epithelium, including *KRT5/KRT6* (CK5/CK6) as specific markers, and they are enriched in patients harboring *TP53* and *BRCA1* mutations ^51^. Basal-like breast cancers with a high expression of growth factor genes (basal-like 2 type) have a particularly poor prognosis and do not respond to chemotherapy ^52,53^ similar to the therapy-induced basal-like subtype we observe in our EAC model.

Since *KRT5, KRT6,* and *KRT16* are not typical keratins for EAC, their expression has not been systematically investigated in EAC to our knowledge. To clarify the clinical role of the therapy-induced basal-like subtype of EAC, we used KRT6 as a marker since KRT6A and KRT6B were among the strongest expression differences in Eso26 RCT vs. control (**Supplementary Fig. 3**). We screened our well-annotated cohort of 728 EAC for the expression of KRT6 and observed a remarkable frequency of 25.7% of tumors expressing KRT6 at detectable levels, 9.6% at high levels, where >25% of tumor cells are positive. The high-level expression was significantly associated with shorter overall survival. Strikingly, this effect was driven by EAC with neoadjuvant RCT or perioperative chemotherapy (**Fig. 6E**) suggesting that the difference in overall survival was to a large extent due to poor treatment response. It is important to note that we determined the KRT6 expression post-therapy which captures the therapy-induced upregulation of KRT6. Patients who did not receive neoadjuvant/perioperative treatment showed a similar trend but did not reach significance. Here, the analyzed samples have not been treated, and it is unknown which of them would have responded with KRT6 upregulation upon therapy. Our findings indicate that every tenth EAC patient has reduced benefit from RCT/chemotherapy. Therefore, it will be important for risk classification to test post-RCT EAC tissue for KRT6 expression. Patients with therapy-induced high KRT6 expression might require more intense surveillance programs or additional therapy modalities, such as immune therapies. It will be important to find new treatment strategies for patients with EAC of therapy-induced basal-like subtype, a group of patients who poorly respond to the current standard of care.

A metaplastic response to radiation of normal squamous epithelium towards stratified squamous epithelium has already been noticed in 1950 ^54^ emphasizing that the transdifferentiation of epithelial cells as a stress response is a widespread phenomenon. A prominent example of the clinical relevance of lineage plasticity is the transdifferentiation of non-small cell lung cancer and prostate cancer to neuroendocrine disease under targeted therapy ^55^. This transdifferentiation poses a highly relevant clinical problem as targeted treatment fails, requiring changed treatment regimens.

Overall, our study revealed a mechanism of adaptation of EAC to RCT that involves the upregulation of stress response keratins and a basal-like subtype. This phenotype overlaps with a response to reduced *BRCA2* function and is associated with short overall survival of treated patients. Future studies will have to clarify the molecular mode of action by which this cell state results in resistance to pave the way for a better treatment of these patients.

## Materials and Methods

### Animal studies

Eight weeks old male nude mice (Rj:NMRI-Foxn1nu/nu) purchased from Janvier labs (Saint Berthevin Cedex, France) were anesthetized with isoflurane for controlled injection of cells. Human xenografts were generated by subcutaneous injection of 5*10^6^ human EAC cells in 100 µl PBS bilateral into both flanks of nude mice. Tumor volume was measured by calliper and calculated with the formula V = π/6 × length × width^2^. Mice were randomly distributed into groups of treatment or control. Treatment was started when a xenograft reached a volume of up to 500 mm^3^. Mice were treated with a combination of chemotherapy and radiation (radiochemotherapy, RCT). At the Department of Radiotherapy and Cyberknife Center of the University Hospital Cologne, radiation was applied locally to the tumor regions in 2 fraction of 11.5 Gy each with a distance of 1 week, yielding in an equivalent dose in 2 Gy fractions of 41.2 Gy assuming α/β =10 Gy in the tumor tissue. Simultaneously with the first radiation dose, chemotherapy was started with i.p. injections of 50 mg/kg/body weight carboplatin twice per week and 20 mg/kg/body weight paclitaxel once per week. Total treatment time was 3 weeks. Mice were sacrificed when one tumor reached a maximum volume of 1000 mm^3^. Close monitoring of mice behaviour did not show any signs of side effects of the treatment. H&E staining of liver, spleen and kidney was performed for all sacrificed mice, with no findings. All animal experiments were approved by the local Ethics Committee of Animal experiments.

### Tissue culture

Human EAC cell lines Eso26, OE19 and OE33 were obtained from Deutsche Sammlung von Mikroorganismen und Zellkulturen (DSMZ, Germany). All cell lines were cultivated in RPMI-1640 (Life technologies, USA) supplemented with 10 % fetal bovine serum (Capricorn Scientific, Germany), 1 % penicillin (Life technologies, USA) and 1 % streptomycin (Life technologies, USA) in a humidified atmosphere of 5 % CO_2_ at 37 °C. All cultures were tested for contamination of mycoplasms by qualitative PCR.

### Immunofluorescence staining of yH2AX

EAC cell lines were seeded on glass slides and allowed to attach for at least 24 hours under optimal culture conditions. Cells were then treated with 1 mM carboplatin in culture media for 24h. Cells were washed twice with PBS, fixed with 2% PFA for 20 minutes and permeabilized with 0.1% TX100 in PBS for 10 min. Blocking was done with 1% BSA in PBS for at least 30 minutes, followed by incubation of primary Phospho-Histone H2A.X Ser139 (20E3, Cell Signaling Technology) in 1% BSA for over-night and subsequently incubation of secondary antibody Alexa-Fluor 488 (#A11034, Thermo Fisher) for 1h.

### Migration assay

EAC cell lines were seeded into multiwell plates and grown to confluency. Cells were starved without serum for 24 hours before a scratch was applied manually with a pipette tip. Cells were washed and again cultured with starving media while migration of cells into the scratch was monitored.

### *In vitro* radio-chemotherapy

EAC cells were seeded into 96-well plates at a density of 2500 cells/well. After 24h, cells were treated with a combination of carboplatin and paclitaxel dissolved in growth media. Right after the start of the chemotherapy, cells were irradiated with 2 Gy (Biobeam GM 8000, Eckert & Ziegler, Gamma-Service Medical GmbH, Germany). Control cells were mock treated. Cells were allowed to grow for 72 h before cell viability was assessed via CellTiter-Glo® 2.0 Cell Viability Assay (Promega, Germany) according to the manufacturer’s instruction and measurement with a Centro LB 960 Microplate Luminometer (Berthold Technologies GmbH & Co. KG, Germany).

### CRISPR/Cas9 editing

EAC cell lines Eso26, OE19 and OE33x were transfected with *BRCA2* Double Nickase Plasmid (sc-400700-NIC, Santa Cruz Biotechnology, USA) targeting human *BRCA2* gene according to the manufacturer’s instruction. Transfected cells were selected by flow sorting for GFP+ cells using a LE-MA900FP flow sorter (Sony) and with puromycin, respectively, and subsequently seeded at a calculated density of 0.5 cells/well for the establishment of single cell clones. Single cells were expanded and DNA was extracted (NucleoSpin Tissue, Mini kit for DNA from cells and tissue, Macherey-Nagel, Germany). With primers flanking the gRNA target site, the extracted DNA was PCR-amplified and analyzed via Sanger sequencing to determine CRISPR/Cas9 edited cells. PCR products from cells identified with a heterozygous sequencing result at the expected CRISPR/Cas9 target site were further analyzed via TOPO® TA Cloning® Kit for Sequencing (Thermo Fisher Scientific, USA) according to the manufacturer’s instructions. Individual PCR-product derived E. coli clones were picked and plasmid inserts were analyzed by Sanger sequencing to determine the spectrum of alleles at the *BRCA2* locus within a single cell colony.

### Biochemical assays

For immunoblot analysis, cells were lysed in Laemmli sample buffer (Bio-Rad, USA) and heated at 95 °C for 5 minutes. Sample proteins were separated via SDS-PAGE and transferred to a methanol activated PVDF membrane via wet transfer blotting. Endogenous BRCA2 was detected with primary antibody anti-BRCA2 (#10741, Cell Signaling Technology, USA) diluted 1:1000 in 5% bovine serum albumin (BSA) in Tris-buffered saline (TBS) containing 0.05% Tween-20.

### Tumor growth rate analysis

The measurements of tumor volume were taken at discrete and unevenly spaced points in time. In order to determine growth rates, we determined derivatives with respect to time from these discrete and noisy measurements, which required smoothing the original data. Suitable tools for smoothing and the estimation of the derivative are a Gaussian smoothing kernel (Gaussian filtering) and a Gaussian derivative kernel, respectively. For smoothing the time series of data points (volume measurements) *x_i_* at times *t_i_* (with time points labelled i = 1, 2,3,… ), we computed the discrete approximation of the Gaussian convolution

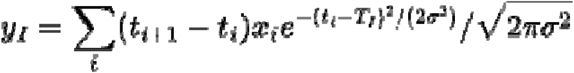

giving smoothed estimates *y_I_* at arbitrary time points *T_I_*. The corresponding estimate of the derivative of the time series follows from the Gaussian derivative kernel

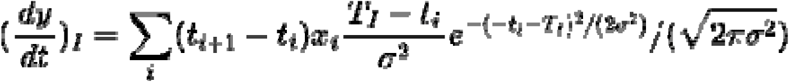

with the estimate of the growth rate at a particular time given by 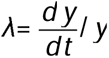 following from the growth equation 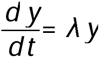. We set σ= 5 days throughout; different choices do not substantially affect the results.

To estimate the growth rate over a period of time, we computed the average of

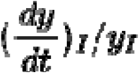

over a specific time interval (**Supplementary Fig. 6**). For time series without treatment, we took the average over the first 15 days of measurement, which corresponds to the exponential growth phase.

Treatment took place over a period of 21 days. To compute growth rates before treatment, we took the average over the days starting with the first non-zero measurement and ending with the start of the treatment. To compute growth rates after treatment, we averaged over the period beginning at day 40 after the start of treatment (19 days after the end of treatment) and ending with the end of the time series. This particular starting point has been chosen to ensure that the growth rate has reached a post-treatment plateau. The figure shows a representative example.

### Xenograft preparation

Xenografts dissected from sacrificed mice were snap-frozen in liquid nitrogen and stored at - 80°C until further processing. When tumor size allowed, a representative tumor region was preserved in 4 % PFA, embedded in paraffin and processed for hematoxylin and eosin staining. The frozen parts of the tumor were ground in mortars under constant cooling with liquid nitrogen to homogenize the tumor material. The ground tissue was split into two cryotubes and stored at -80 °C until further processing. Material from one cryotube was used to extract DNA and RNA using the AllPrep DNA/RNA/miRNA Universal Kit (Qiagen, Germany) according to the manufacturer’s instruction. The second cryotube was used for extraction and in solution digestion of proteins at the Proteomics Core Facility Cologne according to the core facility’s protocol.

### Proteomics

For liquid chromatography mass spectomotry (LCMS) data independent acquisition (DIA), samples were analyzed on a Q Exactive Exploris 480 (Thermo Scientific) mass spectrometer that was coupled to an EASY nLC 1200 (Thermo Scientific). Peptides were loaded with solvent A (0.1% formic acid in water) onto an in-house packed analytical column (30 cm — 75 µm I.D., filled with 2.7 µm Poroshell EC120 C18, Agilent). Peptides were chromatographically separated at a constant flow rate of 300 nL/min and the following gradient: 4-30% solvent B (0.1% formic acid in 80 % acetonitrile) within 74.0 min, 30-55% solvent B within 8.0 min and 55-95% solvent B within 2.0 min, followed by a 6 min column wash with 95% solvent B. These high pressure (HP)LC settings were used for library generation and analytical runs.

For spectrum library generation by gas phase fractionation, aliquots from all samples were pooled and used for spectrum library generation by narrow window DIA of si× 100 m/z gas phase fractions (GPF) covering the range from 400 m/z to 1000 m/z ^56^. The MS was operated in DIA mode. MS1 scans of the respective 100 m/z gas phase fraction were acquired at 60.000 resolution. Maximum injection time was set to 60 msec and the AGC target to 100% (1E6). DIA scans ranging from 300 m/z to 1800 m/z were acquired in 25 staggered 4 m/z windows resulting in nominal 2 m/z windows after demultiplexing. MS2 settings were 15.000 resolution, 60 msec maximum injection time and an AGC target of 1000% (1E6). All scans were stored as centroid.

For data independent acquisition, MS1 scans were acquired from 390 m/z to 1010 m/z at 15k. Maximum injection time was set to 25 msec and the AGC target to 100% (1e6). DIA windows covered the precursor range from 400 - 1000 m/z and were acquired in 75 x 8 m/z staggered mode resulting in nominal 4 m/z windows after demultiplexing. MS2 scans ranged from 300 m/z to 1800 m/z and were acquired at 15 k resolution with a maximum injection time of 22 msec and an AGC target of 100% (1E6). All scans were stored as centroid.

For data preprocessing, Thermo raw files were demultiplexed and transformed to mzML files using the msconvert module in Proteowizard. For spectral library searching, all DIA files were processed using DIA-NN 1.7.16 ^57^. Mouse and a Human canonical Swissprot fasta file were converted to Prosit csv upload files with the convert tool in encyclopedia 0.9.0 ^58^. The following settings were applied: Trypsin, up to 1 missed cleavage, range 396 m/z – 1004 m/z, charge states 2+ and 3+, default charge state 3 and NCE 33. The csv files were uploaded to the Prosit webserver to generate spectrum libraries in generic text format ^59^. Human and mouse spectrum libraries were merged, redundant elution groups were removed and the library was annotated using a merged human and mouse fasta file. A calibrated assays library was generated by searching the predicted library with the 6 GPF runs. Finally, the calibrated assay library was searched with 56 samples runs using double-pass mode and Rt-dependent cross run normalization. Precursor and fragment m/z ranges were set to match the acquisition method. DIA-NN default settings were used for all other parameters.

The DIA-NN output file was filtered for global precursor Q value <0.01 and global protein group Q value <0.01. Identifier columns and protein group MaxLFQ intensities were extracted, transformed to a data matrix and imported to Perseus ^60^. The data set was split into four subsets containing (i) all, (II) human plus shared, (iii) human only and (iv) mouse only protein groups. Each subset was filtered for at least 70% valid values in at least one condition and remaining missing data were imputed using MinDet (q = 0.01) from the imputeLCMD package integrated in Perseus.

### RNA-sequencing

Libraries for RNA sequencing were prepared using the QuantSeq 30 mRNA-Seq Library Prep Kit FWD for Illumina (Lexogen GmbH, Vienna, Austria) according to the low-input protocol. Libraries were sequenced on a NovaSeq 6000 (Illumina) by 1x 50 bases.Downloaded raw data files were first processed with Xenome tool ^61^ to remove host-derived contamination. The remaining, xenograft-only data were mapped to the GRCh38 genome reference using nf-core/rnaseq pipeline version 3.8.1 ^62^. Raw expression read counts obtained as a result were used for differential expression analysis was conducted with DESeq2 ^63^, identifying differentially expressed genes (DEGs). Genes with an absolute log_2_-foldChange value of at least 1 and an adjusted p-value < 0.05 were considered significantly differentially expressed.

### Gene set enrichment analysis

Gene expression data was preprocessed to remove low-expression genes with count per million (CPM) value < 0.5. After preprocessing, the reduced gene expression matrix was utilized for Gene Set Enrichment Analysis (GSEA). The analysis was conducted using the GSEA software version 4.3.3 developed by the Broad Institute, following the standard procedure as described by Subramanian and colleagues ^64^. The genes were ranked according to the signal-to-noise ratio, which was calculated based on the expression difference between conditions. The ranked list was then used as input for the enrichment analysis with gene sets obtained from the Molecular Signatures Database (MSigDB, version 2022.1), specifically the REACTOME collection. Gene sets containing fewer than 15 genes or more than 500 genes were excluded. Permutation test was performed by randomly shuffling the gene labels 1,000 times to generate a null distribution of Enrichment Score (ES). Data normalization was performed using the “meandiv” method resulting in Normalized Enrichment Score (NES). Additionally, the false discovery rate (FDR) was calculated to correct for multiple hypothesis testing, providing an estimate of the proportion of false positives among the gene sets identified as significant. Gene sets with an FDR q-value < 0.25 were considered significantly enriched.

### Whole exome sequencing (WES)

WES was performed using the SureSelect Human All Exon V6 (Agilent) for enrichment according to the manufacturer’s recommendations and were sequenced 2×100 on a NovaSeq 6000 (Illumina). To differentiate between mouse DNA (host) and human xenograft DNA, we utilized the Xenome tool (^61^, version 1.0.0). Only reads classified as graft (graft.fa) were retained for further analysis. Graft.fa reads were aligned to the reference genome using the BWA-MEM ^65^, integrated in the ICGC-Argo pipeline ^66^. For copy number variation (CNV) calling, we employed the Genome Analysis Toolkit (GATK) CNV calling tool (^67^, version gatk-package-4.1.3.0) with Processing Intervals: “gatk CollectReadCounts” and Calling Reads: “gatk CollectReadCounts”. A panel of normals (PON) was created from the control (CTRL) samples of each cell line to serve as a baseline for CNV detection. The CNV calling was then performed using the GATK CNV calling tool, with the PON integrated into the analysis to improve the accuracy of the detected variations. For single nucleotide variation (SNV) calling, a similar approach was used to create a panel of normals. The SNV calling was executed using a modified version of the ICGC Argo pipeline (version gatk-mutect2-variant-calling-4.1.8.0-6.0), tailored in-house to accommodate the analysis of non-matched tumor-normal samples. Variants that were present in both treated and control cell samples were removed to avoid artifacts and pre-existing mutations in the control samples.

### EAC tissue micro array immunohistochemistry and survival analysis

The procedure of tissue microarray (TMA) analysis by immunohistochemistry (IHC) and the cohort of EAC patients have been described earlier ^68^. In brief, tumor specimen were included of 1,161 patients with esophageal adenocarcinoma that underwent Ivor-Lewis esophagectomy with curative intention in between 1998 and 2019 at the University Hospital of Cologne. The local ethics committees approved the study (ethics committee number: 21-1146) and it was conducted in accordance with the declaration of Helsinki. TMA was performed as described elsewhere ^69^. IHC was conducted to assess expression of CK6 (KRT6) with antibody Anti-Cytokeratin 6 antibody [SN71-07] (HUABIO, USA) according to the manufacturer on a automatic staining system Leica BOND-MAX with Leica Bond Polymer Refine Detection Kit (Leica Biosystems, Wetzlar, Germany). For 728 tumors, CK6 IHC was evaluable. CK6 stainings were classified according to their percentage of stained tumor cells (0%= negative, >0-25%= low positive, >25% high positive). Analyses were performed with R version 4.3.0. Overall survival was defined as time between surgery and death or loss of follow-up. P values below 0.05 were considered statistically significant. Chi-square test was conducted to compare qualitative variables. Survival analyses were conducted using Kaplan-Meier curves with respective log-rank tests.

## Supporting information

Suppplementary Table 1

Suppplementary Table 2

Suppplementary Table 3

Suppplementary Table 4

Suppplementary Table 5

Suppplementary Table 6

## Acknowledgements

This study was funded by the Deutsche Forschungsgemeinschaft (DFG, German Research Foundation) through CRC 1310/1 and 1310/2 C04, project number 418074181 and INST 216/1063-1 FUGG (project number 446411360).

## Resource table

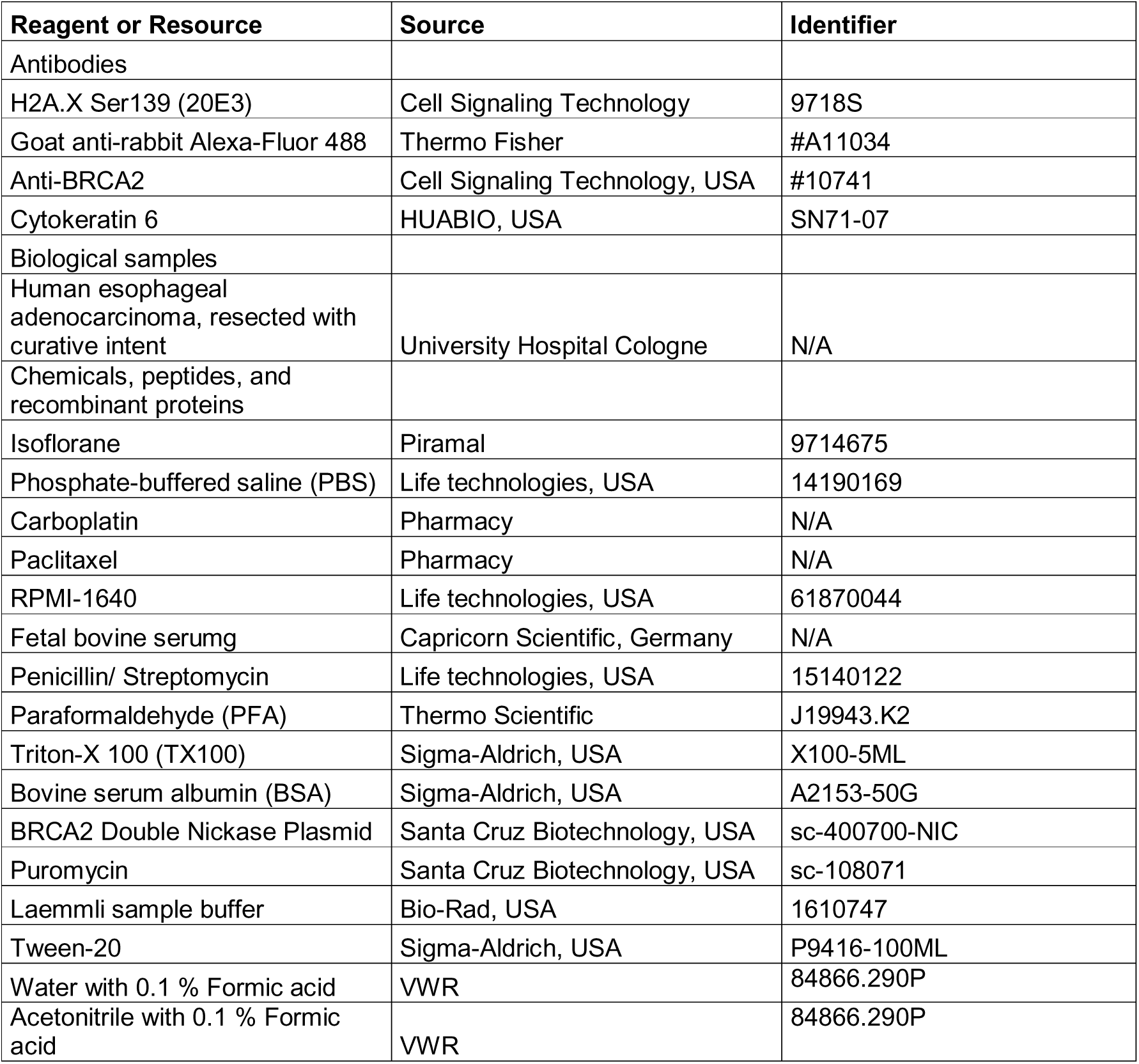
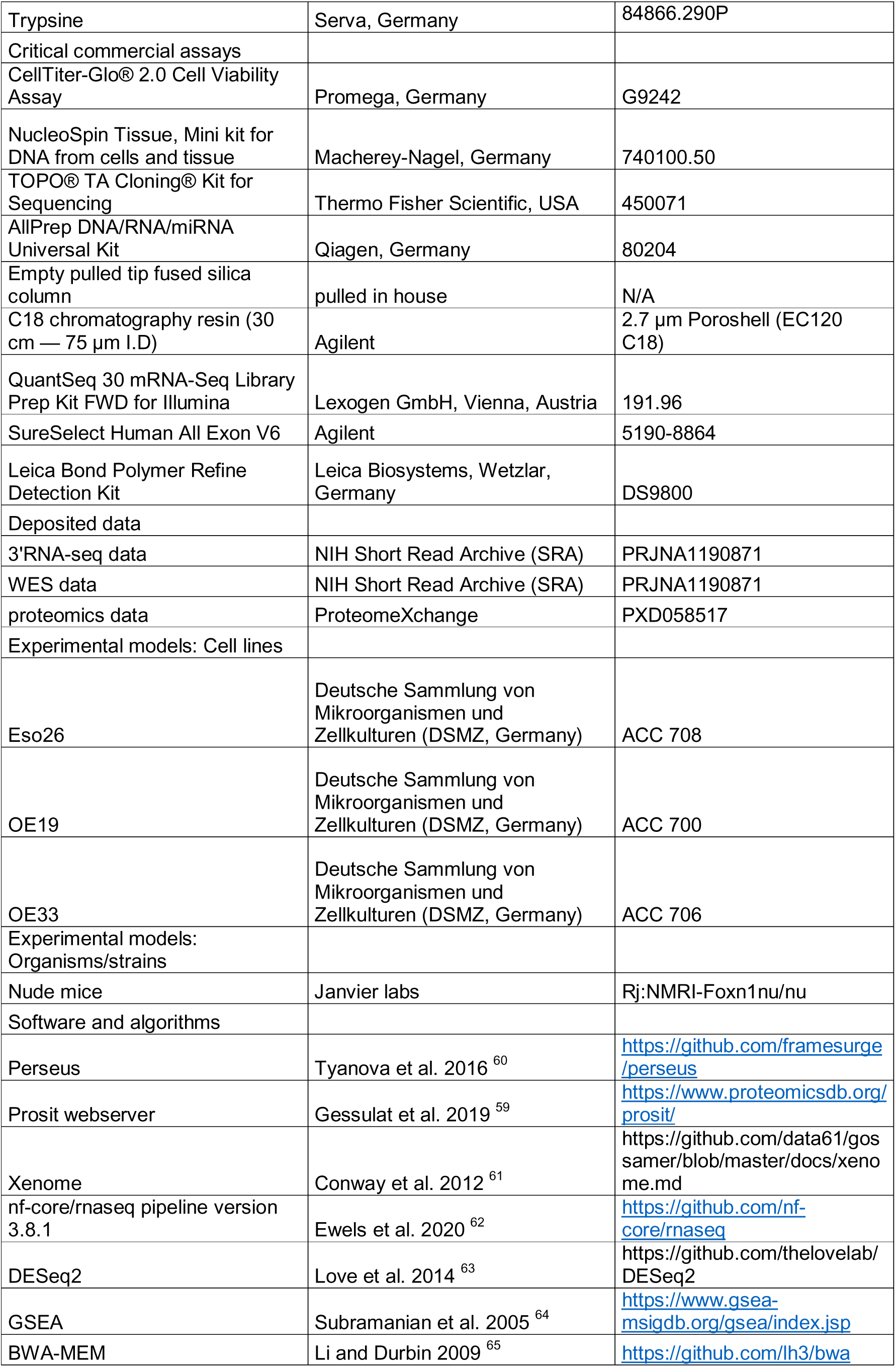
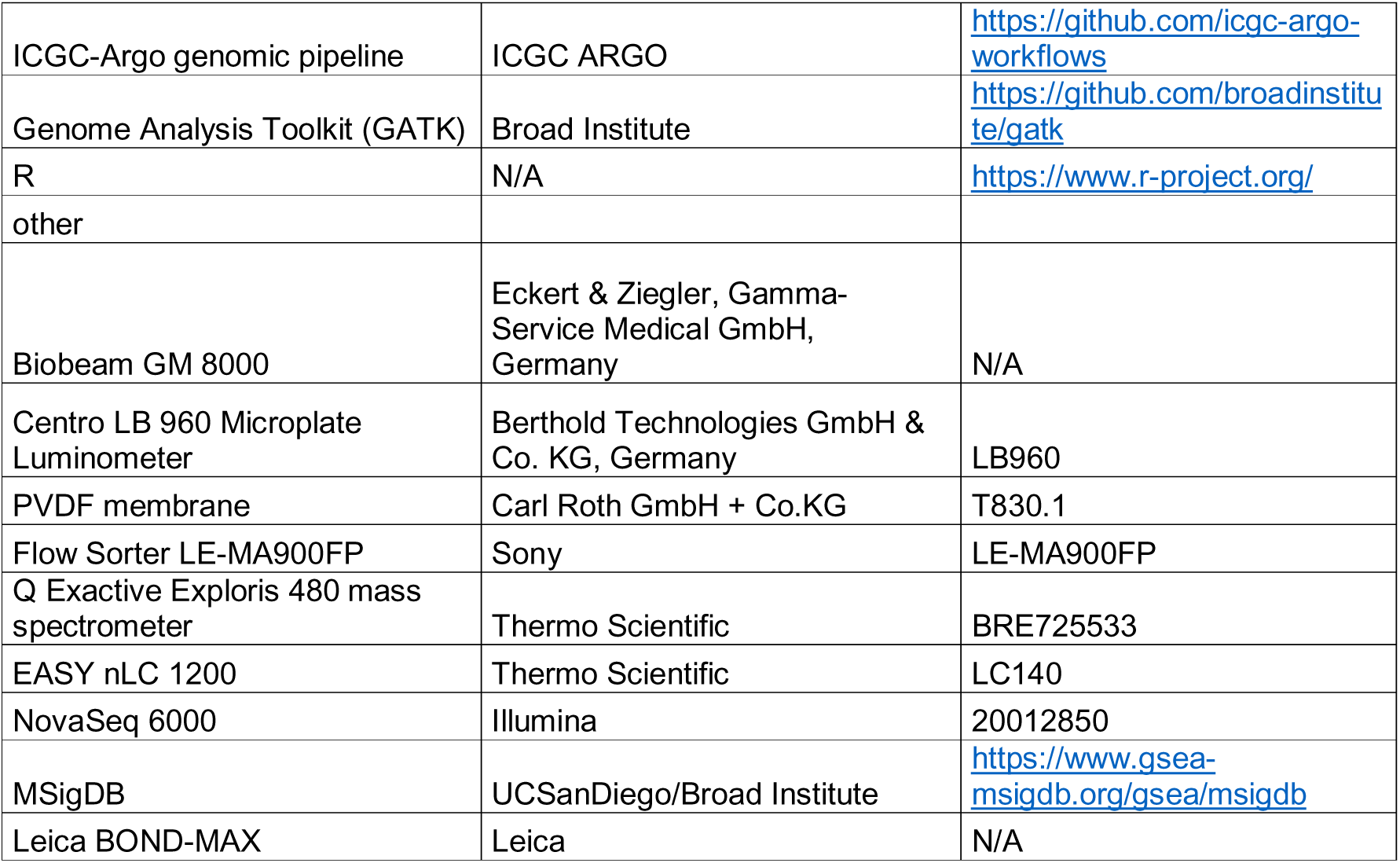

