## Supplementary Material for "Esophageal adenocarcinoma relapse after chemoradiation is dominated by a basal-like subtype"

### Supplementary Information

### Contents

**Supplementary Tables** .......... 2

**Supplementary Figures** .......... 3

### Supplementary Tables

Supplementary Tables are provided as separate data files

**Supplementary Table 1:** Genomic copy number segments of EAC xenografts derived from WES

**Supplementary Table 2:** Small genomic variants when compared to parental control cell line xenograft; *BRCA2*kd after RCT was additionally filtered for variants observed in *BRCA2*kd control

**Supplementary Table 3:** Gene-level transcriptomic data of EAC xenografts of expressed genes derived from 3'RNA-seq measured in counts per million (CPM); averages of CPM per experimental group per gene are displayed

**Supplementary Table 4:** Gene-level proteomic data of EAC xenografts; averages of values per experimental group per protein are displayed

**Supplementary Table 5:** Differentially expressed genes based on 3'RNA-seq and all xenografts of the indicated groups. #P-adj<0.05 indicates the number of significant log2FoldChanges per gene.

**Supplementary Table 6:** Correlation analysis of 3'RNA-seq data of all xenografts of the study for KRT6A (a), KRT6B (b), overlapping genes among the top 500 genes correlating with KRT6A and KRT6B with positive (c) and negative correlation (d)

### Supplementary Figures

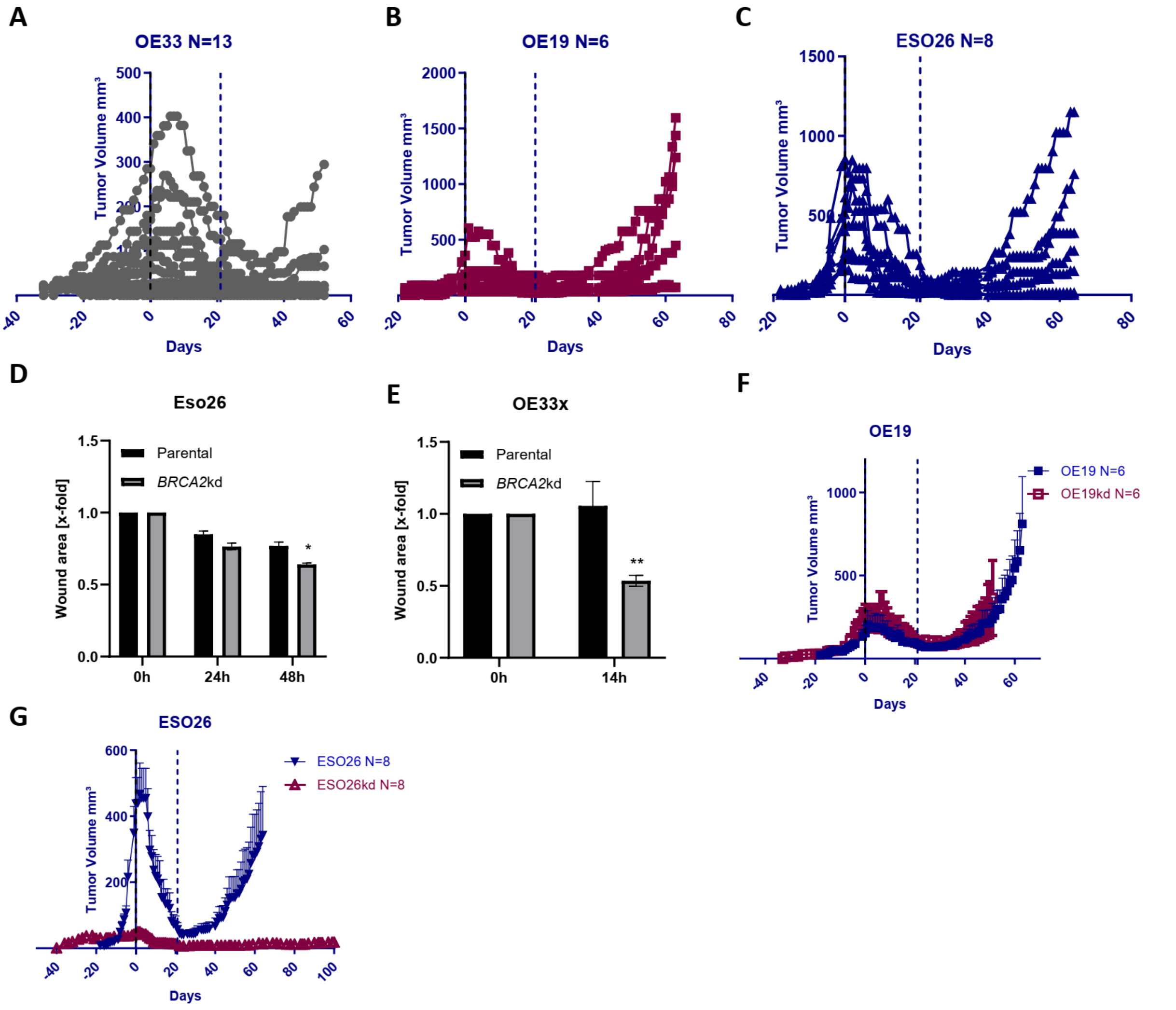


**Supplementary Figure 1:** Individual growth curves of OE33x, OE19, and Eso26 models, migration rates of Eso26 and OE33x models, and cumulative growth curves of *BRCA2*kd models. **(A, B)** Data highlights the distinct progression patterns of tumors. Growth curves demonstrate progression in **(A)** 4 out of 13 tumors in the OE33x model, **(B)** 4 out of 6 tumors in the OE19 model, and **(C)** in 6 of 8 tumors in the Eso26 model following RCT treatment**. (D, E)** *BRCA2*kd cells exhibit enhanced migration abilities in Eso26 **(D)** and OE33x **(E)** -derived xenograft models, as assessed through wound-healing (scratch) assay, in comparison to parental cells (* p <0.05). Reduced relative scratch area represents higher migration rate. One representative of three independent experiments shown. **(F, G)** RCT treatment response tested in OE19 and Eso26 *BRCA2*kd EAC models, respectively. Growth curves for the OE19 **(F)** and Eso26 **(G)** *BRCA2*kd tumor models, showing the initial response to RCT treatment and subsequent post-treatment progression.

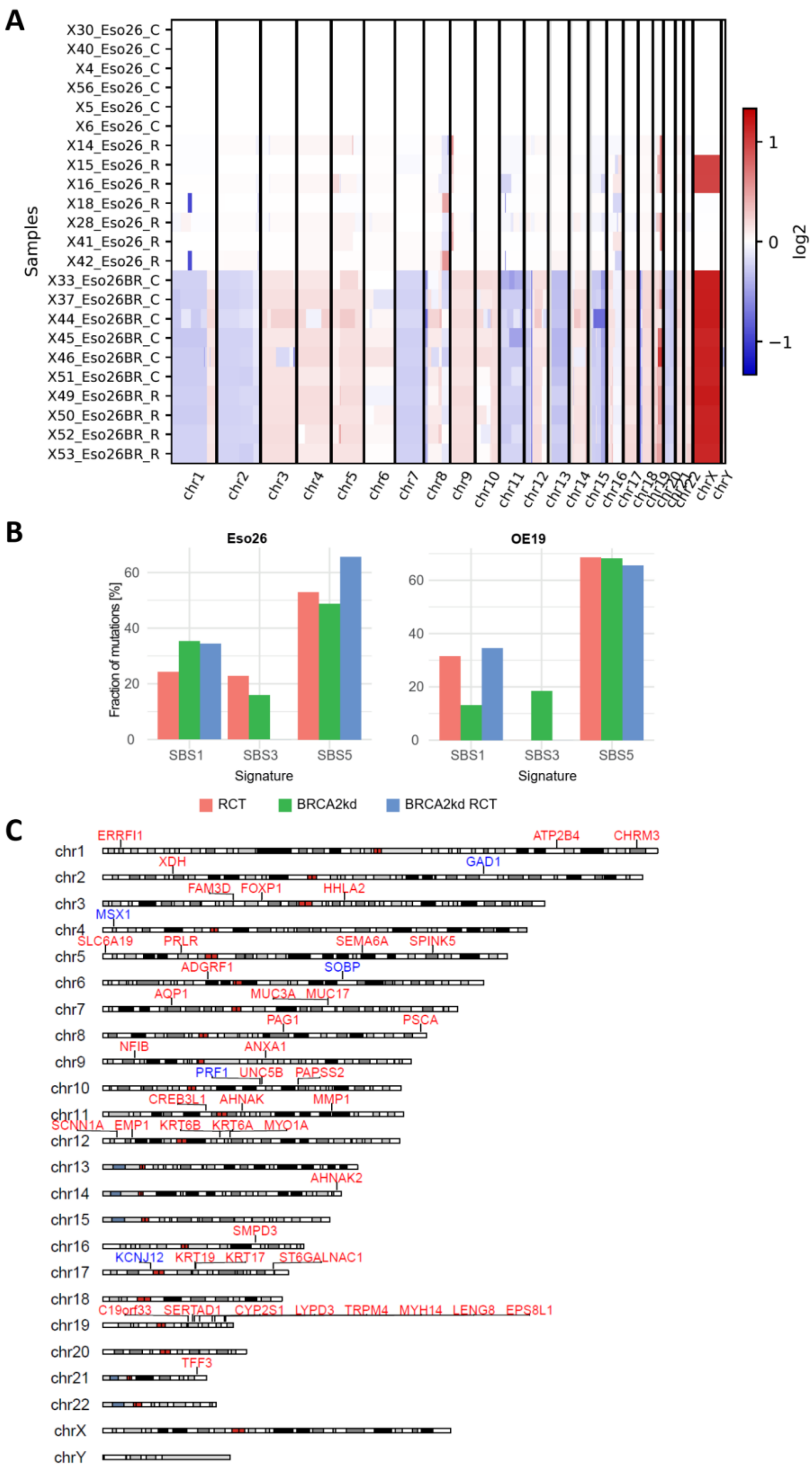


**Supplementary Figure 2:** Genomic changes in Eso26 xenografts. **(A)** Genomic copy number changes induced by RCT and/or *BRCA2*kd treatment in Eso26 xenografts. SCNA genomic heatmap illustrating copy number gain (red). **(B)** Contribution of mutation signatures to the landscape of acquired mutations of Eso26 and OE19 when compared to their parental counterparts. Y-axis indicates the fraction of mutations assigned to the respective mutation signature of all samples of a given experimental group. **(C)** Genomic mapping of the RCT-resistance signature genes on chromosomal bands. 17% of the RCT-resistance genes are located on 19q.

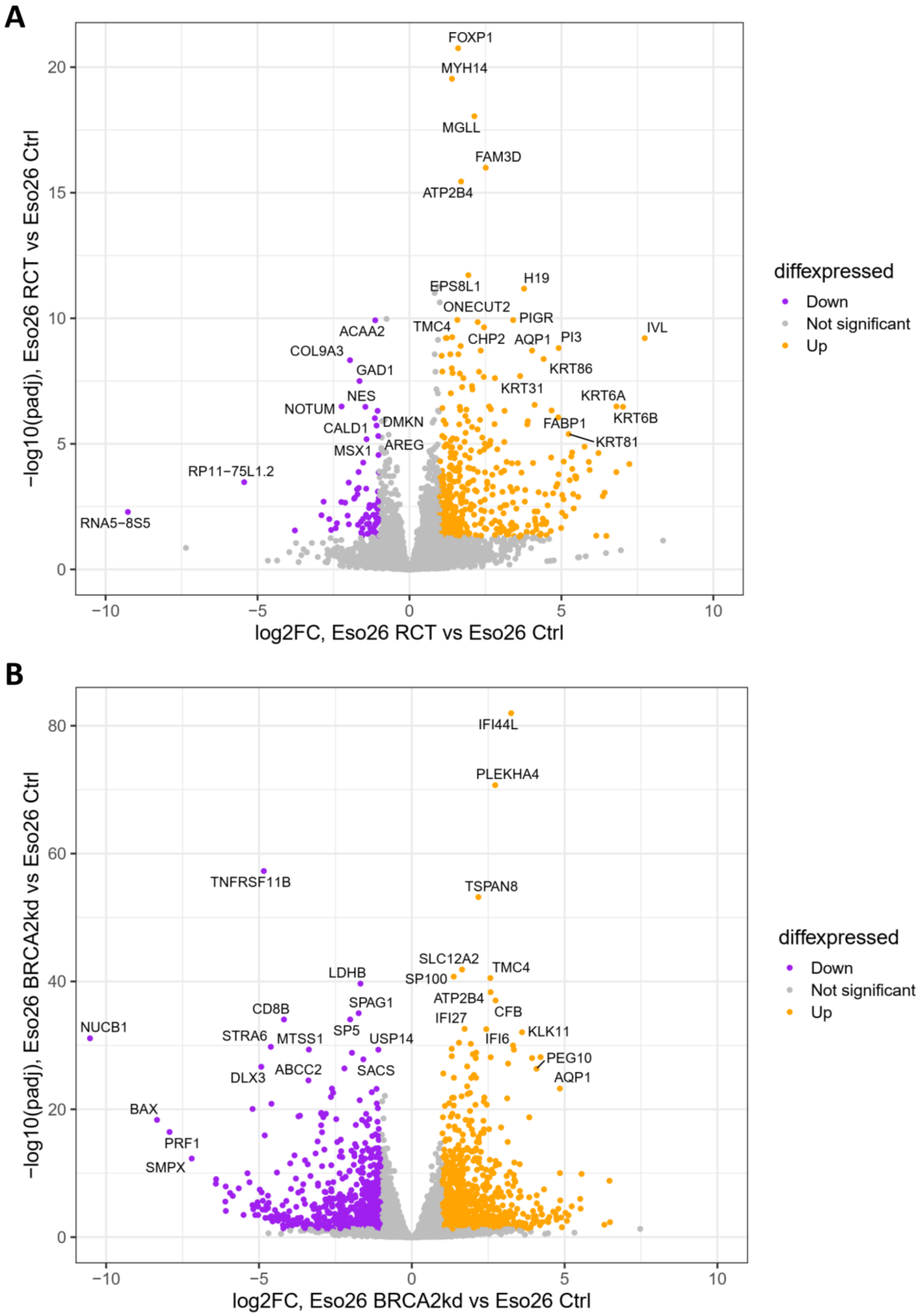


**Supplementary Figure 3:** Differential gene expression analysis of Eso26 xenograft models. 3’RNA-seq of Eso26 xenograft was used for differential gene expression analysis illustrated by volcano plots. (**A**) Eso26 parental after RCT (upregulated in orange top right, downregulated in purple top left) compared to Eso26 parental. (**B**) Eso26 *BRCA2*kd compared to Eso26 parental is shown.

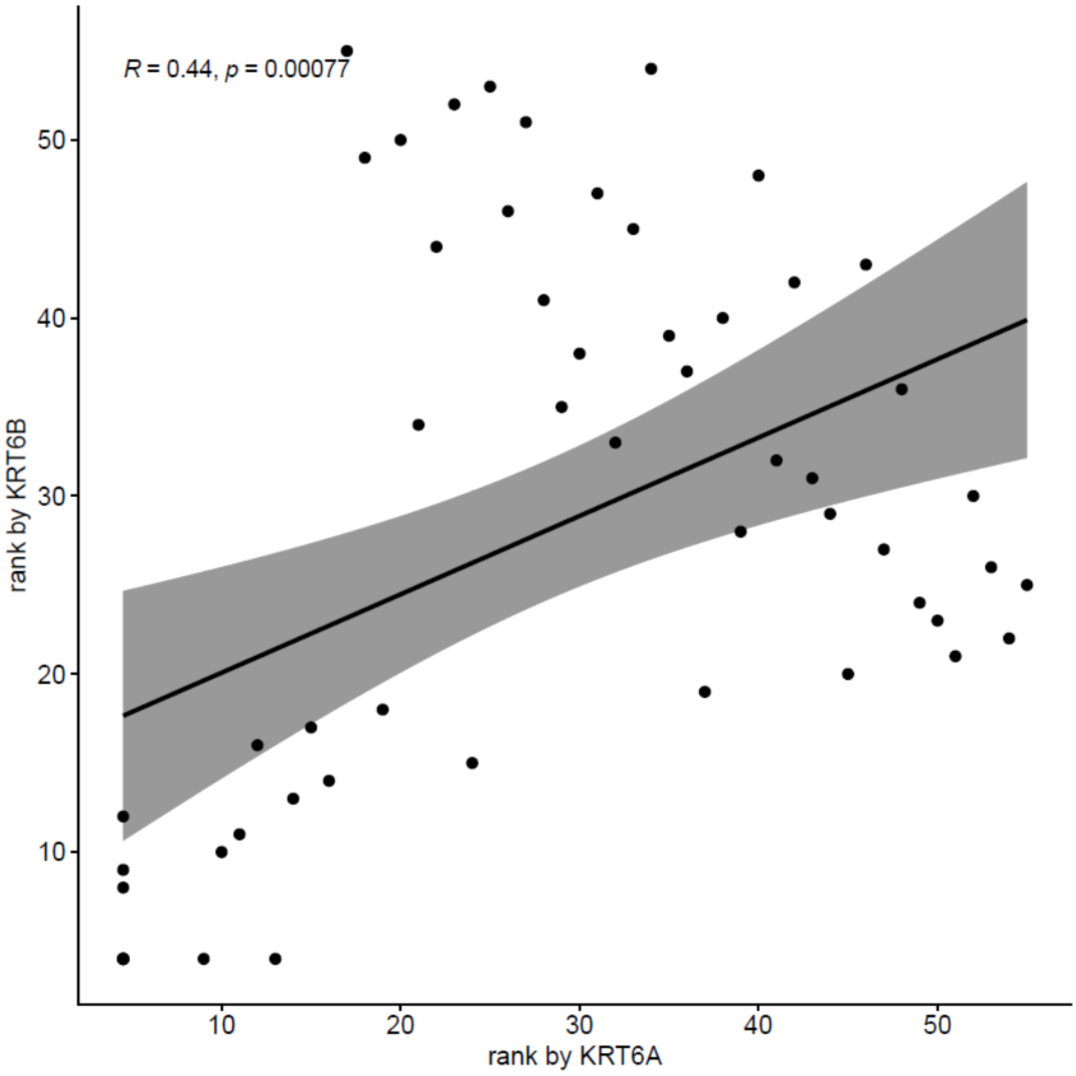


**Supplementary Figure 4:** Correlation analysis of *KRT6A* and *KRT6B* across all xenografts.

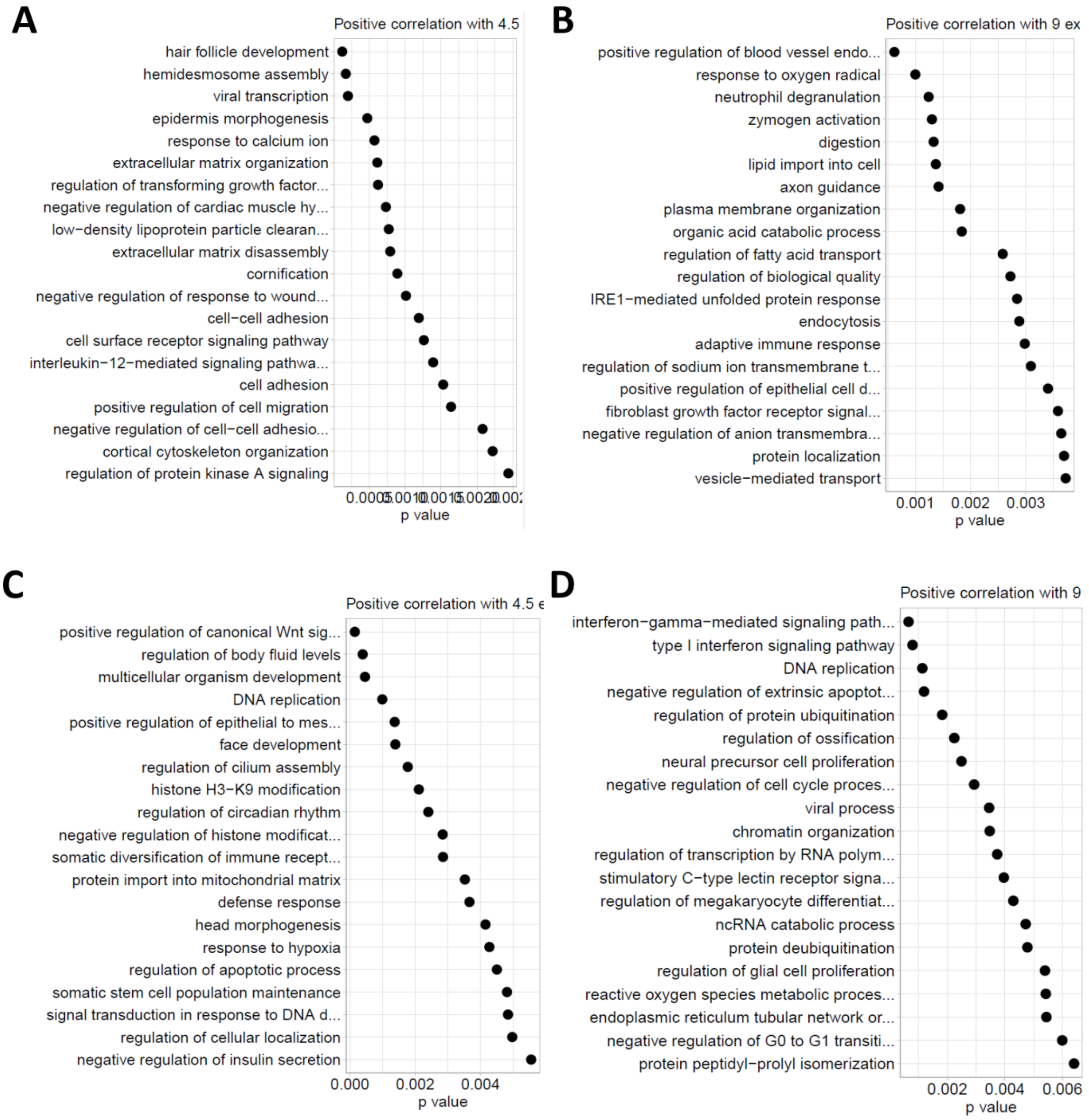


**Supplementary Figure 5:** Gene ontology (GO) analysis of genes correlating with *KRT6* RNA-expression. 3'-RNA-seq data is basis for analysis. All xenografts of the project were used for correlation analysis. GO analysis of biological processes of positively correlated genes with *KRT6A* **(A)** and *KRT6B* **(B)**. GO analysis of biological processes of negatively correlated genes with *KRT6A* **(C)** and *KRT6B* **(D)**. GO terms significantly overrepresented in correlated genes are shown.

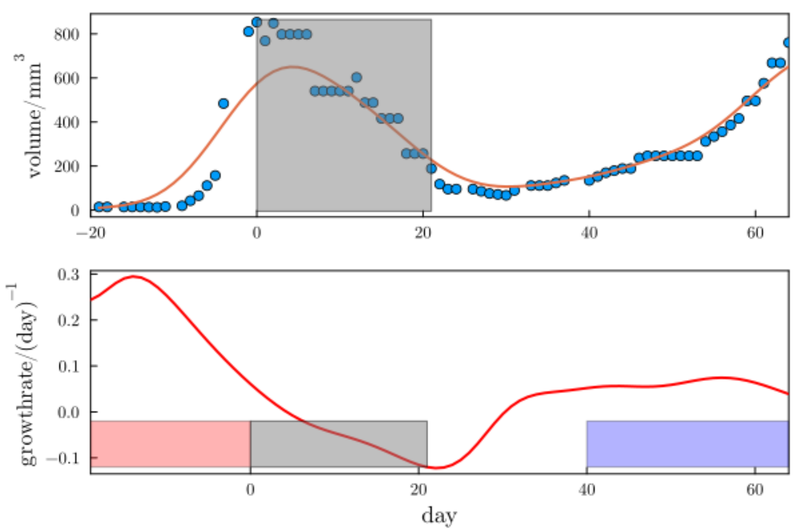


**Supplementary Figure 6:** Smoothed growth curve and growth rates of a representative time series (Eso26_7 from Eso26_parental_interp). The top panel shows volume measurements (blue circles), and the red line the result of Gaussian kernel smoothing. The bottom panel shows the corresponding growth rate estimate over time. The treatment period is indicated in gray, the period over which the rate prior to treatment is averaged is shown in red, the period over which the rate post-treatment is averaged in blue, see text.
